# Resolving antibody avidity through nanoscale antigen patterning

**DOI:** 10.64898/2026.04.18.719169

**Authors:** Iris Rocamonde-Lago, Ieva Berzina, Simon K. Dahlberg, Ian T. Hoffecker, Björn Högberg

## Abstract

Multivalent interactions are fundamental to many biological systems, including how antibodies bind to their respective antigens. Still, their accumulated binding strength, i.e. avidity, resulting from binding and rebinding events of multiple interacting points, remains difficult to measure. Classical assay platforms like ELISA and SPR lack control over antigen positioning, limiting the resolution of the binding characteristics that shape biological outcomes at nanoscale. Here, we present PANMAP, a planar and plate-based assay inspired by ELISA that uses DNA origami to present antigens at defined nanoscale patterns, enabling direct measurements of spatially resolved binding events, termed avidity profiles. Using IgG antibodies as a model system, PANMAP distinguishes between monovalent and bivalent binding states by combining equilibrium absorbance measurements with a simple biophysical model. The avidity profiles reveal how the balance shifts between monovalent and bivalent interactions as a function of antibody concentration and antigen spacing, with intermediate separation distances favoring bivalent binding events and excluded at both near and far distances. This spatial profiling allows decoupling of affinity dependency from avidity profiles to reveal how spatial constraints influence binding equilibria. Our approach fills a longstanding gap in multivalent interaction measurement and offers a new tool for antibody engineering, development of multivalent reagents, therapeutic screening, and mechanistic immunology.

## Introduction

Multivalency occurs when a molecule simultaneously engages multiple sites on its target. It is a common phenomenon in biology and influences the binding behavior of pathogens, cells and molecules, including proteins and DNA, in cellular microdomains, by increasing the strength and specificity of interactions (Ehrlich 1979; Kiessling et al. 2008; Orcutt-Jahns et al. 2023) and modulating the response type (Gestwicki et al. 2002). In the mammalian immune system, the Y-shaped proteins of the immunoglobulin family, commonly called antibodies, are most known for exhibiting multivalency (bivalency) via their two antigen-binding domains. This bivalent binding enhances the affinity (avidity) of a single antibody, like IgG, and prolongs residence time, improving response to pathogens and other external elements (Vauquelin & Charlton 2013). Avidity also enables surface saturation at lower antibody concentrations, facilitating pathogen neutralization and downstream processing by activating the complement system (Kishore & Reid 2000) or the phagocytic Fc receptors (Mammen et al. 1998). Antibodies have also been observed to exhibit bivalent binding to symmetric and repetitive viral or bacterial epitope patterns (Emini et al. 1983; Thouvenin et al. 1997), and digestion of the full-length antibody into monovalent Fab fragments reduces antibody neutralization efficiency (Icenogle et al. 1983). Thus, accounting for bivalency and multivalency is important in therapeutic antibody design (Oostindie et al. 2022), vaccine development (Ku et al. 2022; Rujas et al. 2021), drug development (Buckley et al. 2023) and delivery (Scheepers et al. 2020), and in the understanding biological processes such as autoimmune diseases and pathogen-host responses (Kim et al. 2018).

The bivalent binding potential of an antibody depends on its spatial tolerance, i.e., the physical constraints set by its own native structure and spatial reach. Thus, the two main elements that determine the type of binding are the length of the Fab arms, conserved in most antibodies, and the length and the flexibility of the hinge region, which varies between isotypes and subclasses, and depends on the specificity and environmental conditions. In a typical IgG1 structure, the distance between extended binding sites is around 15 nm (**Figure 1C**), with some cases reported of exceptionally long or protruding paratopes, enabling the antibodies to bridge antigens separated by up to 17 nm (Zhou et al. 2007). Besides the isotype differences, these structural characteristics can be altered or engineered in tailored recombinant antibodies (Kang et al. 2024; Shaw et al. 2019), and are therefore not universal to all immunoglobulins relevant in research or therapy. Hence, antibody binding behavior is highly dependent on the spatial arrangement of antigens (Preiner et al. 2014).

**Figure 1.**
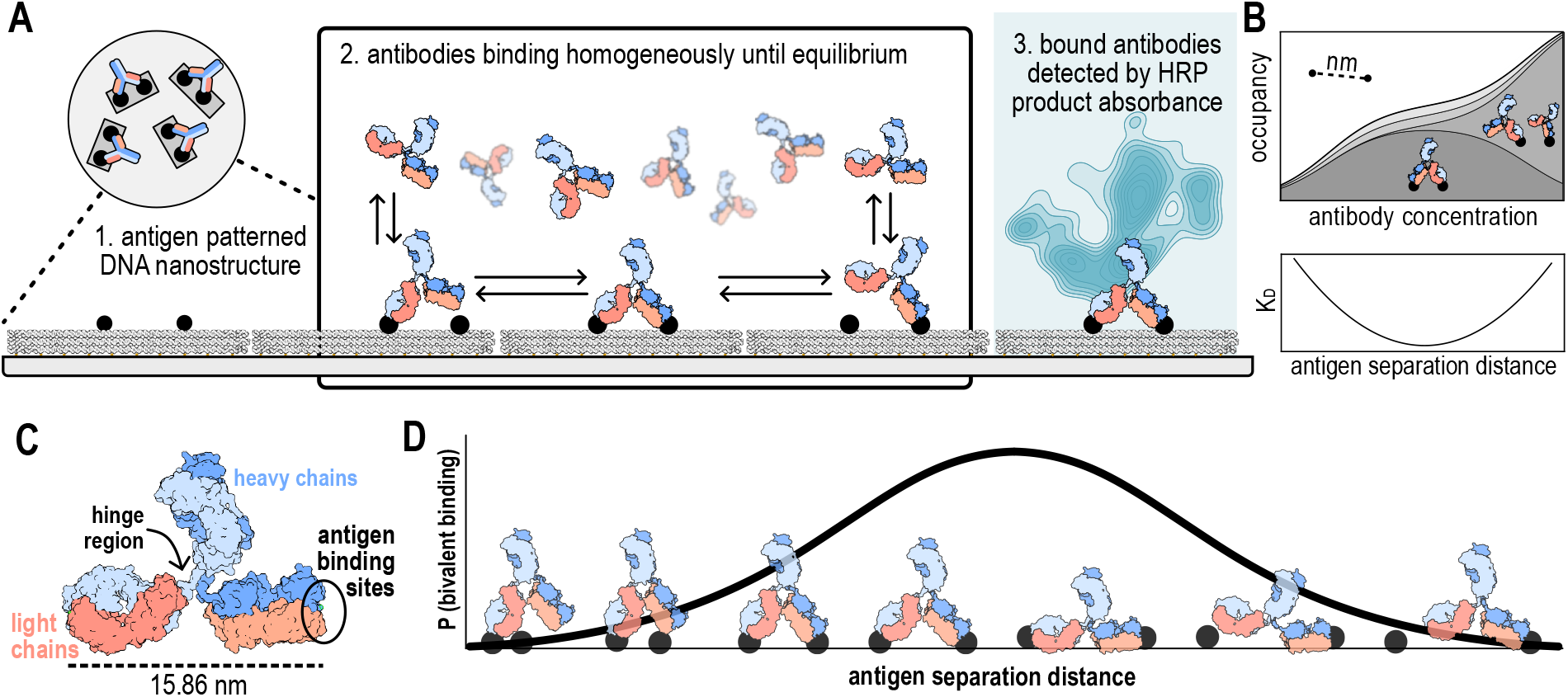
Patterned Antigen Nanostructure-derived Multivalent Avidity Profiling (PANMAP). **(A)** Schematic representation of the PANMAP pipeline. 1) the surface of a plate is coated with the antigen-patterned nanostructures resulting in a homogenous distribution of the antigen. 2) HRP-conjugated antibodies against the antigen of interest bind to the nanopatterns until equilibrium is reached. Thanks to the controlled antigen spatial distribution, the binding mode of the antibodies is also homogeneous. 3) The bound antibodies are detected by measuring the absorbance of the HRP product. **(B)** Output of the PANMAP pipeline: a detailed breakdown of constituent binding states comprising the ensemble as a function of antibody solution concentrations (top), and the affinity dependent on antigen separation distance (bottom). **(C)** ChimeraX illustration of an IgG antibody. PDB ID: 1IGT. **(D)** Schematic representation of binding state progression with antigen separation distance increase.

If antigens are spaced extremely close to one another, a single binder may reach beyond a neighboring antigen, and if multiple binders are present, they may compete for a single binding site causing steric hindrance among antibodies. Conversely, bivalent binding will not occur under conditions where antigens are spaced far apart, beyond the distance that the antibody Fab arms can bridge. Antigen spatial arrangements at the nanoscale are ubiquitous in nature, including highly ordered and repetitive protein arrays on viral capsids (Fang et al., 2022; Kaufmann et al., 2011; Nandhagopal et al., 2002; Sarker et al., 2016), bacterial S-layers (Messner et al., 2008), and membrane-bound protein clusters and multimers (Hartman et al., 2011; Yu et al., 2020; Zhao et al., 2013), providing natural templates for multivalent interactions. Antigen geometry influences B cell activation, in which clustering of antigens leads to clustering of receptors and signal amplification (Dintzis et al. 1976), as well as immune complex formation in which multiple antigens may be cross-linked by multiple antibodies prior to phagocytosis. Consequently, accounting for multivalence for a given antibody-antigen pair requires a combined understanding of the spatial distribution of antigens and the structural constraints of the antibody.

Despite the important role that antigen geometrical arrangement plays in determining antibody binding, it remains a largely overlooked aspect of antibody characterization. This has been likely due to the limitations of current planar binding characterization techniques such as enzyme-linked immunosorbent assay (ELISA), biolayer interferometry (BLI), or surface plasmon resonance (SPR), which do not control for spatial distribution at the nanoscale, making avidity characterization challenging. Without a widely accepted benchmarking for multivalency, the standard practice is to report an apparent monovalent antibody affinity where avidity is often conflated with affinity. Leading to discrepancies in antibody performance in complex samples as a result.

Snapshots of multivalent binding can be observed with microscopic techniques that provide nanoscale spatial resolution such as atomic force microscopy (AFM) (Preiner et al. 2014) and cryo-electron microscopy (cryoEM) (Hewat & Blaas 1996; Smith et al. 1993). These techniques provide a closer look at the multivalent binding system interactions at the individual molecular level. However, they are often low throughput, demand highly specialized instrumentation, and require time and expertise for sample preparation and data analysis.

Spatially controlled ensemble techniques are a third route seeking to bridge the gap between bulk ensemble measurements and nanoscale information. Shaw *et al*. previously showed how nanoscale antigen patterning could be paired with real time antibody binding monitoring by SPR, enabling analysis of multivalent binding kinetics (Hoffecker et al. 2022; Shaw et al. 2019). While this approach provides insight into binding dynamics, extracting steady state behavior from kinetic data requires real-time instrumentation and assumption of rate constants. Direct measurement of equilibrium binding can thus offer a simpler, more accessible, and standardized alternative, both experimentally and for data interpretation.

Here, we developed a simple planar and plate-based method for directly measuring equilibrium binding under conditions of controlled antigen spatial separation, Patterned Antigen Nanostructure-derived Multivalent Avidity Profiling (PANMAP), which produces a quantitative representation of the avidity between an antigen-antibody pair (**Figure 1A**). Like ELISA, PANMAP is a colorimetric assay to measure antibody binding, but it positions antigens in a precise and controlled pattern on the surface of DNA nanostructures. These are made by the DNA origami method for the self-assembly of three-dimensional DNA nanostructures, based on the hybridization of short oligonucleotides (staples) to their complementary regions in a long circular DNA molecule (scaffold) (Benson et al., 2015; Douglas et al., 2009; Rothemund et al., 2006). The addressability of DNA origami allows for antigen spatial arrangement with nanoscale precision. Thus, PANMAP maps the functional relationship between spatial separation of antigens and the equilibrium binding behavior of an antibody.

In this work, we establish an assay system where the concentration of antibodies in solution is known, spatial separation between single antigens is controlled, and the total number of DNA nanostructures and bound antibodies is measurable. This enables a detailed mapping of constituent binding states between the binding molecules and their antigens. The resulting avidity profile of an interacting pair is composed of a set of model parameters that may be determined in a single experiment. Thus, we examine the ensemble of steady state binding of bivalent antibodies on 2-antigen-patterned DNA origami nanostructures over varying separation distances between individual antigen sites (**Figure 1B**). With PANMAP, the characterization of bivalent antibody interactions can be standardized to inform the design of antibody therapeutics and vaccine development and to provide a higher level of understanding of antibody-antigen binding in a closer setup to their biological context.

## Results and discussion

We sought to establish a simple plate-based assay that could precisely reveal avidity effects in antibody analytes using standard laboratory equipment. We aimed to resolve the functional connection between antigen spatial separation distance and the steady state binding characteristics of an IgG-type antibody. To achieve this, we used DNA origami nanostructures to control the spatial arrangement of a small molecule hapten target, coupled with a colorimetric readout system, enabling us to measure the quantity of bound antibodies at equilibrium. We examined multiple experimental separation distances over an interval from 0 to 35 nm, and multiple antibody concentrations in order to construct a mapping between observable measurement and constituent binding states (**Figure 1**).

### Spatially defined antigen libraries exhibit multivalency

We first designed a DNA origami-based system to control the spatial arrangement of antigens. We assembled a library of DNA nanostructures as molecular breadboards using oligonucleotides that partially protrude from an array of possible positions for antigen anchoring (**Figure S1**). The DNA origamis have a rod-like shape (Hoffecker et al. 2022; Shaw et al. 2014; Shaw et al. 2019; Smyrlaki et al. 2024; Spratt et al. 2024), and the anchoring nucleotides can have varying degrees of rotational freedom to facilitate or restrict the interaction between the antigens and the antibodies against them. In a classic ELISA experiment, the planar surface of a microtiter plate is coated with antigens that are unspecifically adsorbed to the plastic or to a coated surface, creating a heterogenous distribution of targets and their orientations. This approach lacks spatial control of antigen immobilization. Thus, the distribution, orientation, and distance between antigens is random, creating uneven binding site densities that result in a complex mixture of antibody binding states. This leads to a mixed readout that corresponds to a composite mix of antibody-antigen interactions. The binding event is reported via any bound antibody, and it is the same regardless of whether the binder is monovalently or bivalently bound. This makes it difficult to assign physical and, particularly, spatial properties appropriately. In contrast, in a PANMAP assay, the antigens are first precisely patterned at the nanoscale on a DNA origami, that is then attached to the surface plate via biotinylated anchors, creating a homogenous orientation of the epitopes on the surface.

We designed structures displaying a range of distances with a small, stepwise, continuous increase of separation between two antigens that would give us a detailed framework of how the antibody binding mode changes. By creating combinations of specific protruding positions, we selected a library of eight separation distances ranging from 4 to 35.7 nm (**Figure S2,S3**) to encompass the IgG range of monovalent and bivalent binding. These distances, however, can be varied to adapt to the structural or relevant characteristics of any binding partner of interest. As our antigen, and for proof-of-concept experiments, we selected the hapten digoxigenin (**Figure S4**) (Lai et al. 2016), due to its widespread use, chemical stability, and small size that helps minimize the confounding effects of orientation, i.e. accessibility to epitope, on antigen binding.

We incorporated the hapten antigen during the nanostructure folding using digoxigenin-modified protruding staples (**Figure S1**). In our DNA origami library, we also included an empty structure without antigens as a background control for nonspecific antibody binding and a 1-digoxigenin origami as a monovalent reference for the evaluation of the 2-digoxigenin structures (**Figure S2**).

First, to validate the folding quality and to verify the true dimensions of the DNA nanopatterns, we employed both negative stain (NS) transmission electron microscopy (TEM) (**Figure 2B**) and cryo-electron microscopy (cryo-EM) (**Figure 2A**). With TEM, we observed predominantly straight DNA rods matching the designed number of helices and overall dimensions. When we reconstructed the nanostructure from cryo-EM data using 3DFlex tools (Punjani et al. 2017), the full-length origami could be resolved to a resolution of less than 10Å, which could be improved down to 7Å when focusing on the core region of the origami, revealing a structure closely matching the expected helical arrangement and overall *in silico* design.

**Figure 2.**
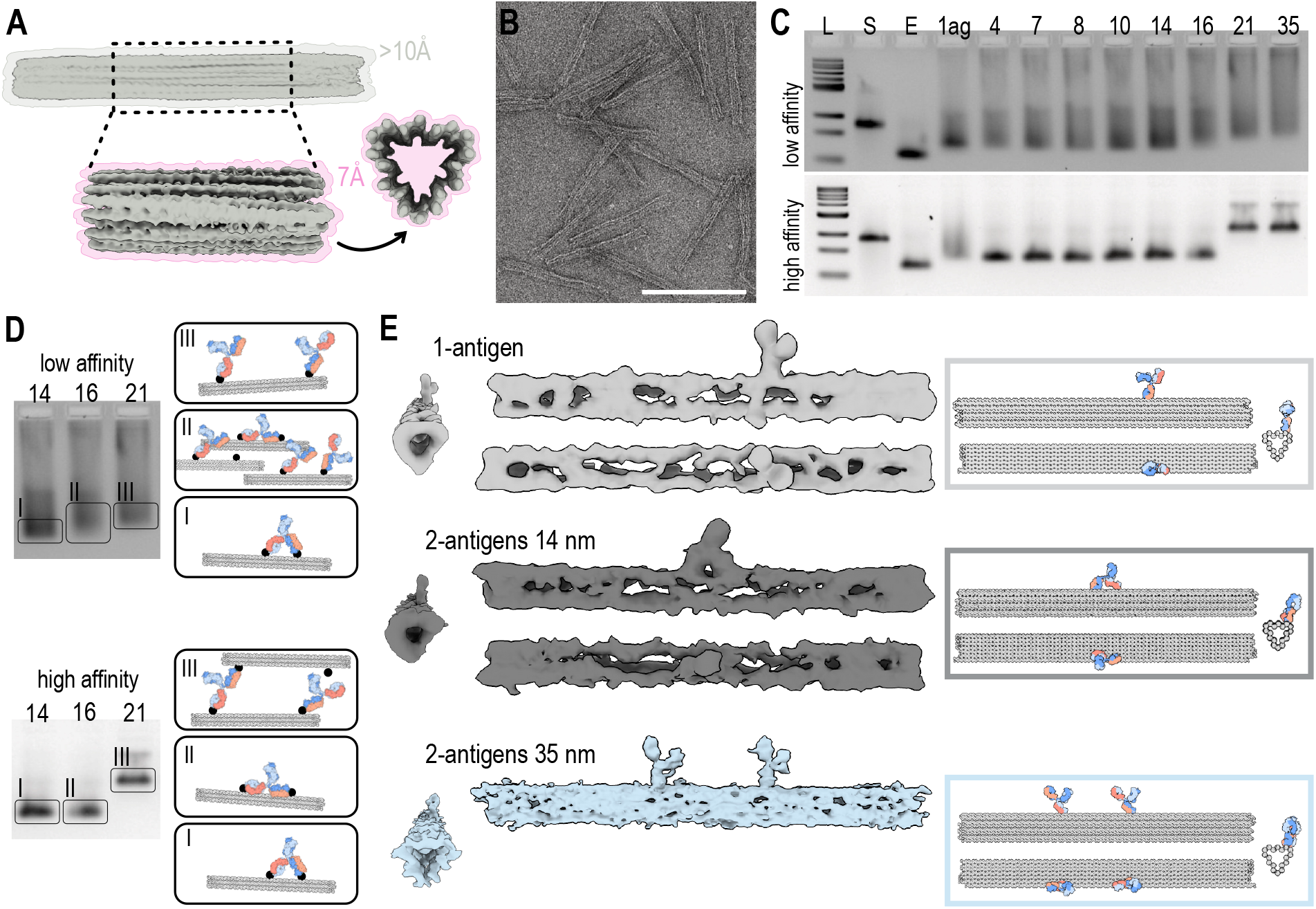
Characterization of the antigen-coated nanopatterns and their interaction with antibodies. **(A)** Cryo-EM density map of the rod DNA origami with corresponding achieved resolutions. **(B)** TEM micrograph showing a field of view of empty DNA nanostructures. Scale bar 140 nm. **(C)** Agarose gel electrophoresis of the antigen-coated nanopatterns after incubation with an excess of low affinity (top) or high affinity (bottom) α-digoxigenin antibodies. L: DNA ladder, S: scaffold, E: empty nanostructure 1ag: 1-antigen nanostructure, 4-35: 2-antigen nanostructures with separations of 4nm, 7nm, 8nm, 10nm, 14nm, 16nm, 21nm and 35nm. **(D)** Representation of the possible antibody states comprising the electrophoretic bands from the gels in (C). 14: 14 nm 2-antigen nanopattern, 16: 16 nm 2-antigen nanopattern, 21: 21 nm 2-antigen nanopattern. **(E)** On the left, TEM 3D class average reconstructions of antibody-bound DNA nanopatterns with 1 antigen (top), 2 antigens separated by 14 nm (middle) or 2 antigens separated by 35 nm (bottom), from different perspectives. ChimeraX illustration of the antibody configurations observed on the TEM three-dimensional reconstructions (right).

To assess the functionality of the antigen-patterned DNA structures for multivalent binding, we incubated the nanostructures in solution with an excess of α-digoxigenin antibodies and then analyzed their migration by electromobility shift assay (EMSA) (**Figure 2C, 2D, Figure S5**). For this experiment, we used two types of α-digoxigenin antibodies, one with high affinity, and another with low/intermediate affinity. The effect of antigen spatial arrangement on antibody multivalency was observed as shifting bands during EMSA. For both binders, there was a marked retardation in migration speed when comparing the empty DNA rod to the 1-antigen structure as a result of a single antibody binding. The nanopatterns with antigens separated by distances ranging from 4 to 14 nm displayed a migration pattern comparable to the structure with 1-antigen, suggesting the presence of a single antibody, bound either monovalently or bivalently.

In the case of the low affinity antibody, the nanopatterns with antigens separated by 21 and 35 nm exhibited a shifted band, which we attributed to the presence of two monovalently bound antibodies (**Figure 2C**). The migration of the 16 nm nanostructure presented a smeared pattern between a fast band similar to the 4-14 nm DNA rods, and a slower band similar to the 21-35 nm DNA rods. In contrast, the high affinity antibody had a sharp band with the 16 nm nanopattern, migrating at the same speed as the 4-14 nm structures, most likely corresponding to the same binding state. Whereas the 21-35 nm nanopatterns showed a notable electrophoretic shift, larger than the one seen for the same samples with the low affinity antibody.

We attribute the distance-dependent shift to a situation in which low affinity antibodies favor a single monovalent or bivalently bound states at short distances. At larger distances of 21 or 35 nm, the predominant state is two monovalently bound antibodies. At intermediate distances (16 nm), a mixture of the two states is apparent. The absence of this mixed condition in the case of the high affinity antibody could be attributed to a greater tolerance for strain due to the greater affinity of each Fab arm, resulting in a sharper transition peak between favorable states (**Figure 2D**). The large shift of the 21 and 35 nm pattern appears to correspond to DNA rods crosslinked by antibodies binding to two inter-structure antigens. Our interpretation of the antibody states from EMSA is shown in **Figure 2D**. This ambiguity further highlights the need for additional constraints to be able to determine the constituent binding states in the ensemble.

To visualize the different binding states favored by different antigen separation distances, we reconstructed negatively stained antibody-bound nanopatterns after imaging them by TEM (**Figure 2E**). We chose three representative examples of configurations studied with EMSA: 1-antigen structure, 14 nm 2-antigens structure, and 35 nm 2-antigens structure. The 2D and 3D reconstructions (**Figure 2E**) confirm that the actual antigen positions matched the designed sites on the DNA rod. The major configurations consisted of a single monovalently bound antibody for the 1-antigen structure, a single antibody engaging both arms at the two designed antigen positions for the 14-nm 2-antigen structure, and two antibodies bound monovalently for the 25-nm 2-antigen structure (**Figure 2E**). These configurations were in agreement with both the EMSA data and the expected maximum distance between binding sites on the IgG1 Fab arms (**Figure 1C**). We illustrated the observed configurations using structural reference models of the DNA rod and antibody (Harris et al. 1997) (**Figure 2E**). These results demonstrate that antigen separation distance influences the prevalence of multivalent binding states. However, the specific composition of monovalent and bivalent binding states, and how these are affected by environmental conditions, is not revealed with these approaches. The spectrum of compositions spans predominantly monovalent binding at shorter distances to a mixture of bivalent states at intermediate and longer distances, indicating an evolution of compositions that could benefit from individual state resolving capability (**Figure 1C**).

### Binding quantification yields equilibrium structure occupancy

We next sought to evaluate PANMAP for quantifying the binding equilibrium of antibodies on the spatially defined antigen-DNA nanostructure library. To do this, we incubated an HRP-labeled antibody with the DNA origami-coated plates until the interaction reached equilibrium, like in an ELISA experiment. As readout of the PANMAP assay, we measured the change in absorbance upon the enzymatic reaction of the HRP-antibody with an added colorimetric substrate. This readout is a semi-quantitative measurement of the total amount of antibody bound to the antigen-patterned nanostructures at the assay endpoint (**Figure S8**). To further convert the absorbance into absolute numbers of bound antibodies, and to obtain fully quantitative information, we experimentally generated a standard curve with known antibody concentrations and extracted a conversion factor from the linear regime of the signal response (see Methods). With this conversion function, we transformed all the PANMAP experimental HRP readouts to antibody amounts at the assay endpoint. These will then be the data used for the binding analysis. Then, we used the converted data of the 1-antigen nanostructure as a reference to estimate the number of immobilized structures per well relative to the saturation point (**Figure S8**). We independently validated the uniformity of nanostructure immobilization across the plate wells using qPCR targeting the scaffold of the DNA nanostructures after the PANMAP assay. This experiment confirmed that the reference-based normalization was sound (see Methods, **Figure S9**).

To determine the contributions of monovalent and bivalent binding to the different antigen separation distances, we further processed the experimental data to obtain what we termed as equilibrium occupancies. The equilibrium occupancy (Φ_*eq*_) describes the average number of antibodies bound per nanostructure at the endpoint of a PANMAP experiment, and it gives integral information about the avidity of the interaction. We obtained the equilibrium occupancies by normalizing the experimental PANMAP data using the calculated quantity of nanopatterns immobilized (**Figure S8**). Thus, we could compute the corresponding equilibrium occupancies of all samples, beginning with the 1-antigen structures, at each of the tested antibody concentrations in solution (**Figure 3A**). This conversion of the experimental data, i.e., HRP signal, and its normalization by number of immobilized nanopatterns enabled us to express the observed antibody binding in absolute molecular terms, laying the basis for resolving the composition of multivalent binding states.

**Figure 3.**
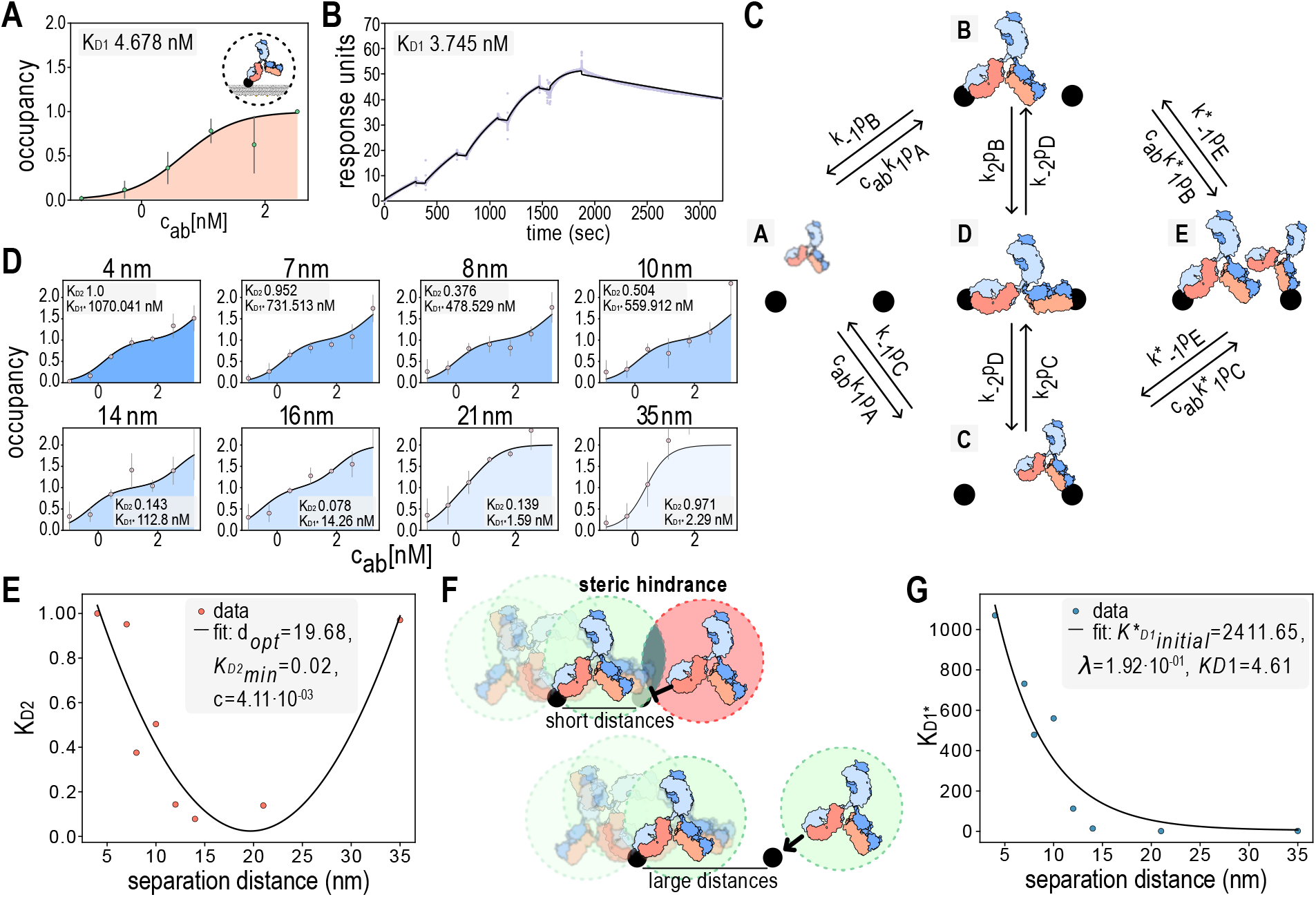
The antibody occupancy for the 1-antigen and 2-antigen nanopatterns. **(A)** Occupancy curve for the 1-antigen nanopattern with the calculated K_D_ in the grey inset. The dots represent the occupancy data obtained experimentally by PANMAP and the black line is the fitted model output. n=3 independent replicates, mean ± S.E.M. **(B)** SPR recording of a single cycle kinetics experiment where the 1-antigen nanopattern was immobilized on the chip and increasing concentrations of HRP-labelled α-digoxigenin antibody were injected (12 nM, 24 nM, 48 nM, 96 nM, 192 nM). The reported dissociation constant K_D_ is in the grey inset. **(C)** Representation of the 5-state Markov model of antibody binding in a 2-antigen system with the binding and unbinding between monovalent and bivalent processes. **(D)** Occupancy curves for the 2-antigens nanopatterns separated by the designated distances above the graphs. The dots represent the occupancy data obtained experimentally by PANMAP and the black line is the fitted model output. The monovalent-to-bivalent dissociation constant (K_D2_) and the sterically hindered monovalent dissociation constant (K*_D1_) for each antigen separation is reported in the grey inset. n=3 independent replicates, mean ± S.E.M. **(E)** Parabolic model of bivalent interconversion constant K_D2_, fitted to experimental PANMAP data as a function of antigen separation distance. **(F)** Illustration of the steric hindrance effect between neighboring antibodies when antigens are spaced in close proximity. **(G)** Fitted exponential decay model of steric hindrance effect on monovalent dissociation K*_D1_. Each datapoint corresponds to the measured K*_D1_ value for a given antigen separation distance.

The resulting PANMAP occupancy profile for the 1-antigen nanopattern showed a strictly monotonic and sigmoidal (single inflection point) approach towards a maximum occupancy of 1 antibody per structure for the antibody highest concentration (**Figure 3A**). From this 1-antigen structure sample, we computed the monovalent affinity of the interaction as a dissociation constant (K_D_) of 4.678 nM. We validated this measurement by SPR, immobilizing the 1-antigen structure and allowing it to bind to increasing concentrations of the same antibody. Using SPR, we obtained a K_D_ of 3.745 nM, in close agreement with the PANMAP measurement, reflecting the reliability of our method to determine monovalent affinities (**Figure 3B**). The 1-antigen nanopattern occupancy provided us with a reference to interpret the PANMAP occupancy signal of all the other 2-antigen structures (**Figure 3D**).

As seen in the **Figure 3D** curves, the shape of the occupancy curves evolved as antigens were separated by greater distances. For the shorter distances, at the highest antibody concentrations in solution, the maximum recorded equilibrium occupancy Φ_*eq*_ was of 1 antibody per structure, whereas a maximum of 2 antibodies was observed for greater separation distances of 16 nm, 21 nm and 35 nm. Interestingly, the occupancy curves from the short and intermediate distances exhibited a distinct doubly sigmoidal (two inflection points) shape characterized by a convex shoulder at intermediate concentrations, distinct from the singly inflected sigmoidal 1-antigen curves. This additional feature smoothed out with the increased antigen separation of 16 nm and completely disappeared by 21 nm (**Figure 3D**). We hypothesized that this shoulder corresponds to a scenario in which favorable bivalent IgG binding represents the main occupancy at low-to-intermediate concentrations. Once the number of antibodies in solution is increased, the competition for binding is incremented, displacing one of the antibody arms and transitioning into a mix of one or two-antibody monovalent states. This disturbance in the bivalently state can be explained by the structure of IgG-like antibodies, where the Fab arms would have a strained configuration when binding close-by antigens, making them more prone to dissociate when a higher amount of competing partners are present. When the antigens are placed at an intermediate spatial separation, matching a more natural configuration for the antibody arms, the bivalent binding is stronger and the transition to monovalent antibodies is less abrupt, as evidenced by the 16 nm curve (**Figure 3D**).

Another interesting feature that we observed from the occupancy curves is that even though at high antibody concentrations they seem to be bound monovalently, only the distances of 21 nm and 35 nm reach a 2-antibody occupancy, with the rest of distances achieving only 1.5 antibodies on average. We interpret this as an effect of the steric hindrance that antibodies can pose to one another when their target antigens are spaced too close together, displacing each other from their binding positions. When the antigens are further away, the antibodies have enough space to both bind monovalently comfortably (**Figure 3F**).

### System constraints reveal composition of valency states

To further understand the evolution of antibody binding as a function of antigen separation distance, we developed a probabilistic model of steady-state antibody binding for a 2-antigen system. This model is defined by its probability to assume one of five possible states: (p_A_) an empty structure, (p_B_) a monovalently occupied structure with one antibody bound to one antigen, (p_C_) a monovalently occupied structure with one antibody bound to the other antigen, (p_D_) a bivalently occupied structure with one antibody bound to both antigens, and (p_E_) a saturated structure with two antibodies bound monovalently to each antigen (**Figure 4A**). We converted this model derived from elementary rate laws governing the transitions between the mentioned binding states into an expression for equilibrium occupancy as a function of the solution phase antibody concentration (**Figure S11)**. In this way, we can fit our observed PANMAP data into the model and obtain even more precise information on the binding state of each combination of antibody concentration and antigen separation measured experimentally. The expression of the model is:

**Figure 4.**
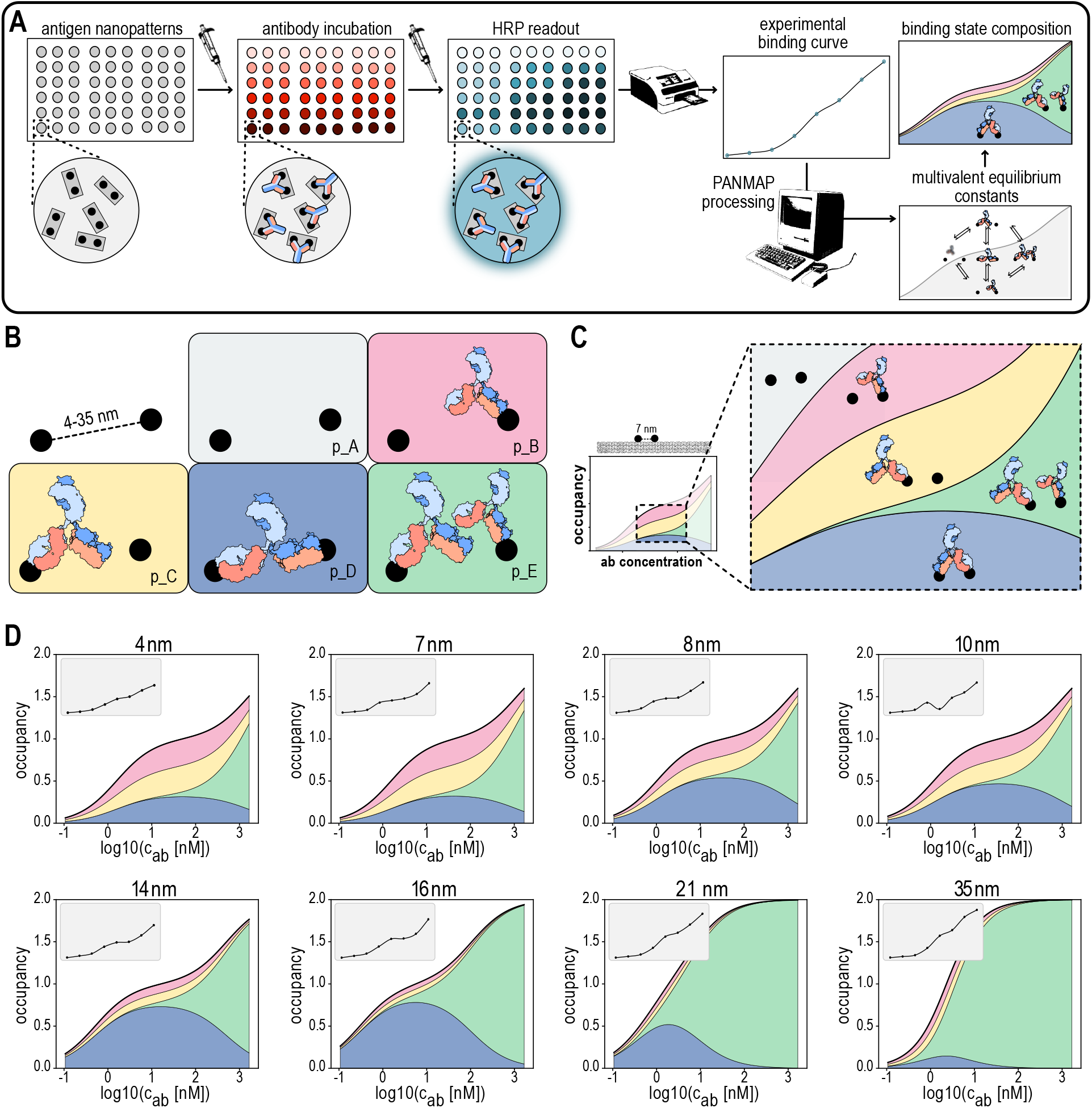
Weighted probability of each state composing the antibody occupancy for antigen-nanopatterns. **(A)** Representation of each possible state in the two-antigen system. The colors of each state correspond to the stratified plots in (B) and include empty (p_A_), monovalent binding (p_B_, p_C_), bivalent binding (p_D_), and two monovalent antibodies (p_E_). **(B)** Weighted probability of the transient states composing the steady-state occupancy for a range of concentrations of antibody in solution for each separation of the two-antigen nanopatterns.

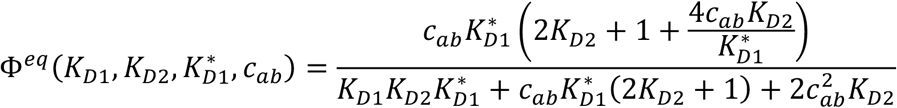

Where *K*_*D*1_ is the monovalent equilibrium dissociation constant, *K*_*D*2_ is an equilibrium dissociation constant for the monovalent to bivalent transition process and 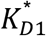 is the monovalent equilibrium dissociation constant for binding when an adjacent antigen is already occupied by a monovalently bound antibody.

We fitted this equilibrium occupancy model to the experimental PANMAP data to obtain the predicted occupancy curves and a corresponding parameter set for each separation distance (**Figure 3D, 4B,D**). Here we again observed a doubly-sigmoidal curve with a convex shoulder present at intermediate solution concentrations predicted by the occupancy model (**Figure S10**), and also predicting that distantly separated structures have a single inflection point (resembling a familiar sigmoidal ELISA curve and our 1-antigen pattern result) whilst more closely spaced antigens had two inflection points.

Whereas a classical affinity measurement (e.g. via an ELISA experiment) aims to describe binding through a single dissociation constant K_D_ (hereafter referred to as K_D1_), binding to patterns as a function of antigen separation distance presents additional degrees of freedom to account for and to inform about the multivalent states. An additional dissociation constant K_D2_ enables us to describe the equilibrium relationship between monovalently- and bivalently-bound states. Likewise, classical affinity measurements, measured at the sparse limit of bound molecules to minimize possible interactions, do not account for the exclusion of multiple antibodies binding to nearby adjacent sites due to steric hindrance, a phenomenon that we would expect to affect the monovalent binding parameter K_D1_ (K_D1*_ for distinction). The PANMAP pipeline enabled us to model the functional relationship between equilibrium binding and spatial antigen separation while incorporating these effects.

For the bivalent K_D2_ values we observed a parabolic progression as a function of distance (**Figure 4E**), with both long and short distances corresponding to more dissociation (weaker conversion to the bivalent state), and intermediate distances displaying the least dissociation (strongest conversion to bivalent state). We phenomenologically modeled this distance dependence or spatial tolerance with a quadratic function:

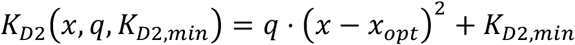

where q is a rigidity factor, representing the sharpness of the parabola, x_opt_ is the optimal bivalent binding distance or minimum of the curve, and K_D2,min_ is the minimum value of K_D2_.

In the case of K_D1*_, we observed a non-linear decline (**Figure 4F**), with short distances corresponding to weaker binding, and longer distances approaching the un-hindered K_D1_. This could be interpreted as reflecting a steric hindrance, i.e. when an existing antibody is monovalently bound to one antigen, at close distances its presence will interfere with the binding of a second antibody. At the limit of well-separated antigens, the two sites will not interact, and the steady state dissociation would approach that of the normal 1-antigen monovalent process (K_D1_). We modeled this dependence with an exponential decay:

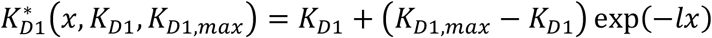

Where l is a steepness parameter indicating the sharpness of the transition and K_D1,max_ is the maximum value of K_D1_ (high dissociation due to hindrance) observed at the limit of no separation between antigens. The model predicts that at large separations, K_D1*_ = K_D1_ since the steric hindrance effect would not take place.

Altogether, the models of K_D1*_ and K_D2_ enable us to describe the steady state binding characteristics of antibodies when two antigens are separated by an arbitrary distance. For a given antibody-antigen system, we may concisely represent a predictive multivalent avidity profile as the following set of parameters: M = {*K*_*D*1_, *q, x*_*opt*_, *K*_*D*2,*min*_, *K*_*D*1,*max*_, *l*}.

Finally, having obtained the dissociation constant parameters by fitting the occupancy signal data, we could then use the model to predict the expected steady-state proportion of each state, allowing stratification of the ensemble occupancy signal into individual contributions (**Figure 4A & B**). This revealed a more detailed picture of the trade-off taking place as both concentration and separation distance are increased (**Figure 4C**). Thus, a greater contribution from monovalent 1-antibody states (monovalent and bivalent) in the close distance ranges (4 nm to 10 nm) and low to intermediate concentrations, and a transition over to saturated 2-antibody states in long distances (16 nm – 35 nm) and high concentrations was observed.

## Discussion and conclusions

In the mammalian immune system, the Y-shaped proteins of the immunoglobulin family rely on multivalent interactions to enhance binding strength, or avidity, and prolong residence time on target surfaces. This multivalent potential is governed by the antibody’s spatial tolerance, defined by the reach of its Fab arms and the flexibility of its hinge region. While these interactions are critical for biological processes like pathogen neutralization and B-cell activation, they remain difficult to characterize using standard platforms like ELISA or SPR, which lack control over antigen positioning at the nanoscale. Consequently, avidity is often conflated with affinity in reported measurements, leading to discrepancies in predicted antibody performance. Although high-resolution techniques like AFM and cryo-EM can be used to observe individual multivalent states, their low throughput and specialized equipment requirements highlight the need for an accessible, ensemble-based method to resolve the functional relationship between antigen spatial distribution and binding equilibrium.

To address the lack of spatial control in traditional binding assays, we developed the planar plate-based assay PANMAP, a platform that improves bulk assays by providing molecular nanoscale resolution of the measured interaction. Exploiting the programmable addressability of DNA origami, we transitioned from the random, heterogeneous antigen distributions of a typical ELISA setup to a system of precisely defined molecular breadboards that is still plate-based and easy to execute without specialized instruments. PANMAP is able to disentangle monovalent and bivalent IgG antibody binding at equilibrium by controlling nanometer-scale spacing between antigens displayed on a DNA origami surface. This method enables equilibrium binding measurement of antibodies under conditions of controlled antigen spatial arrangement and subsequent parameterization of a concise model describing multivalent binding, that we termed avidity profile. Obtaining this avidity profile does not require specialized equipment, with the entire end-to-end procedure consisting of a series of pipetting and absorbance measurement steps, with an entire set of parameters tested from a single plate.

The PANMAP method utilizes rod DNA nanostructures to display antigens, such as the hapten digoxigenin, at precise nanoscale intervals ranging from around 4 to 36 nm. By integrating these patterned structures into a plate-based, colorimetric assay similar to ELISA, the system measures equilibrium antibody binding through an HRP-product absorbance, which is subsequently converted into absolute equilibrium occupancies (the average number of antibodies bound per structure). This approach, validated by SPR, TEM, and cryo-EM, successfully distinguishes between monovalent and bivalent binding states by applying a 5-state biophysical model. The results demonstrate a clear avidity profile where bivalent binding is favored at intermediate distances (reaching an optimum near 16 nm) but is restricted at short distances due to steric hindrance, and at long distances by the physical reach of the antibody arms.

Applying the PANMAP method revealed the trade-offs governing multivalent binding behavior: the antibody bivalent binding spatial tolerance dependent on antigen separation distance, as well as the impact of steric hindrance on monovalent binding. Both of which are also affected by the antibody concentration in solution. Our results for the most common IgG-type binder, provide a rich and quantifiable understanding of multivalent antibody binding. This can be applied to a variety of pipelines, enabling screening, selection, and design of different antigen-antibody pairs for a more effective binding characterization. Our approach makes affinity and avidity available as tuning factors for specificity and tailored for downstream applications.

Part of the innovation of the PANMAP method lies in its dedicated processing pipeline, which translates standard colorimetric readouts into specific molecular binding modes. By establishing a conversion function from an experimental standard curve, the system transforms raw HRP absorbance units into the absolute number of antibodies bound at the assay endpoint. These data are then normalized by the quantity of immobilized DNA nanostructures to calculate the equilibrium occupancy (Φ_*eq*_), representing the average number of antibodies per patterned site. To make this complex biophysical analysis more accessible, we developed a user-friendly pipeline that automates the transition from raw data to a stratified breakdown of the ensemble. By fitting the experimental occupancy curves to a 5-state Markovian model, the software deconstructs the signal into the individual probabilities of a structure being empty, monovalently occupied, bivalently bridged, or saturated with two antibodies. Thus, all the graphs, including the calculation of the equilibrium constants is outputted by only importing the raw data from, e.g. a plate reader. Allowing users to resolve the precise composition of valency states and extract key multivalent equilibrium constants without requiring expertise in rate laws or stochastic modeling.

Beyond its utility as a characterization tool in molecular interactions research, PANMAP offers a unique lens through which to study host-pathogen interactions. For example, it could help elucidate whether specific viral surface patterns force suboptimal monovalent responses, potentially explaining the neutralization escape of certain viruses. Simultaneously, it can also help deepen the understanding of responses to viruses and bacteria by assessing the antibodies produced to them in the context of avidity, to answer questions like: are the patterns in the surface forcing the selection of antibodies that must bind monovalently for an efficient immune response? Furthermore, the relevance of PANMAP extends to the field of therapeutic design and mechanistic immunology. By providing a high-resolution map of spatial tolerance, PANMAP can refine the discovery pipeline for bispecific and recombinant binder drugs, allowing selection of candidates with an optimal geometric fit before advancing to costly cell and animal experiments, or even clinical trials.

The inherent flexibility of DNA origami as a breadboard means that this approach is easily customizable for various molecules and complex patterns (Douglas et al., 2012; Huang et al., 2024; Kwon et al., 2020; Song et al., 2026; Wang et al., 2025; Zhao et al., 2021), making it applicable to multispecific drugs that require the simultaneous engagement of multiple targets, a booming trend in treatment for cancer or hematological diseases (Duell et al., 2019; Klein et al., 2024), and that vary in their nature, including peptides, aptamers, small molecules, or proteins. Ultimately, by making avidity a tunable factor rather than a conflated variable, PANMAP provides the structural insights necessary to engineer the next generation of high-precision multivalent reagents.

## Methods

### DNA origami preparation

#### DNA origami design

The DNA nanostructure was designed using the caDNAno software [http:/_/cadnano.org] following the 3D design parameters described in Douglas *et al*., 2009 (Douglas et al. 2009). The rod structure was based on the origami used by Shaw *et al*., 2014; whose design and structure files can be downloaded from the repository [https://nanobase.org] (Shaw et al. 2014). Briefly, the rod structure is an 18-helix bundle with hexagonal lattice that contains protruding staples on the top and bottom surfaces (**Figure S1**). The bottom protruding staples are modified with biotin for immobilization on the PANMAP plate, or with 21-nucleotide protruding staples for immobilization by DNA complementarity on the SPR experiments. The top protruding staples are modified with the antigen Digoxigenin to create a range of different separation distances. The different antigen combinations are shown in **Figure S2**.

#### Scaffold and staples preparation

The p7560 scaffold was produced from an M13 bacteriophage variant using bacteria as a host. Phage-competent *E. coli* TOP10-were used to inoculate a volume of 50 ml of Lysogeny broth (LB) (Merck) supplemented with 10 μg/ml of doxycycline (Merck, #D9891-1G) and grown overnight in a shaker at 37°C and 250 rpm. In the morning, 35 mL of the overnight culture was used to inoculate 500 mL of 2xYT medium (Merck) supplemented with 5 mM MgCl_2_ and 50 μl of antifoam 204 (Merck, #A8311-50ML). The culture was grown using Ultra Yield flasks (LAB Sweden AB, #931136-B) in a shaker at 37°C and 250 rpm until reaching the optimal density of 0.5 at optical density 600 nm (OD_600_), then the phages were added at a multiple of infection (MOI) of 1 and incubated for another 4 h. To remove the bacteria the culture was centrifuged for 30 min at 4000 rcf and 4°C, with the phage remaining in the supernatant. To precipitate the phage, the supernatant was mixed with 4% PEG 8000 and 0.5 M NaCl for 15 min until the solution becomes cloudy. Then, the phage was pelleted by centrifugation at 4000 rcf for 15 min at 25°C and resuspended in 10 mM Tris 1 mM EDTA pH 8.5. To remove any remaining bacteria, the resuspended phage was further centrifuged at 15000 rcf for 15 min at 4°C. To lyse the phage protein coat, an equal volume of Buffer P2 (Qiagen, #19052) containing 0.2 M NaOH and 1% SDS was added and mixed gently by swirling. Immediately after, the mixture was neutralized by inversions with Buffer P3 (Qiagen, #19053) containing 3M KOAc pH 5.5. The sample was incubated in an ice water bath for 15 minutes before centrifugation at 16500 rcf for 10 min at 4°C. The supernatant containing the ssDNA was transferred to new bottles, and precipitated by mixing with at least 2 volumes of 100% ethanol and incubated on ice for 30 min. The DNA was pelleted by centrifugation at 16500 rcf for 30 min at 4°C and washed with 75% ethanol before a final centrifugation step for 10 min. The ssDNA pellet was airdried and resuspended in 10 mM Tris 1 mM EDTA pH 8.5 and the final concentration was determined by absorbance at 260 nm using a Nanodrop 2000 spectrophotometer (Thermo Fisher Scientific). The scaffold was diluted into 200 nM stocks with 5 mM Tris pH 8.5 and the quality was assessed by agarose gel electrophoresis with the same protocol as for folded nanostructures explained below. The core, edges, and biotinylated staples were purchased from Integrated DNA technologies (IDT), and the digoxigenin-modified staples were sourced from Biomers (**Figure S4**).

#### Folding of patterned DNA origami nanostructures

A typical folding reaction consisted of 20 nM p7560 scaffold, 100 nM staple set, 5 mM Tris, 1 mM EDTA and 8 mM MgCl_2_, final concentrations in 1 ml total volume. The staple sets consisted of 400 nM core and edge staples, 800 nM biotinylated staples for immobilization, and 4800 nM digoxigenin-modified protruding staples, to make a 5-fold, 10-fold and 60-fold molar excess of each respective staple compared to the scaffold during folding. The folding reaction consisted of an initial annealing step at 65°C for 4 min, then a temperature ramp from 65°C to 50°C at 1 min/0.7°C, followed by 50°C to 35°C at 1h/1°C and 20°C for storage until washing the folded structures. The excess staples were removed by PEG precipitation diluting the folding reaction 1:1 with buffer containing 15% PEG 8000 (Thermo Fisher Scientific), 5 mM Tris 1 mM EDTA, and 505 mM NaCl (Merck), and mixing by inversions. The DNA nanostructures were precipitated by centrifugation at maximum speed for 25 min, then the supernatant was discarded and the pellet was resuspended in folding buffer with 20 mM MgCl_2_. The precipitation and resuspension steps were repeated three times, and the last pellet was resuspended in 1x PBS with 5 mM MgCl_2_ by incubating for 30 min at 37°C and 300 rpm. The final concentration of DNA nanostructures was determined by absorbance at 260 nm using a Nanodrop 2000 spectrophotometer (Thermo Fisher Scientific).

### Characterization of DNA origami nanostructures

#### Gel electrophoresis

The folding quality of the DNA nanostructures was assessed by agarose gel electrophoresis. Typically, a final concentration of 8 nM for each sample was loaded onto 2% agarose gels in 0.5x TBE supplemented with 10 mM MgCl_2_ (Merck) and 0.5 mg/ml ethidium bromide (Merck). For the gels with antibody-bound DNA origamis, antigen-patterned structures were incubated with an antibody molar excess of 1000–fold for the 1 antigen and 4 nm to 16 nm samples, or 2000-fold for the 21 nm and 35 nm samples. The experiment was done with a high affinity α-digoxigenin (Thermo Fisher Scientific, #9H27L19) antibody and a low/intermediate affinity α-digoxigenin HRP (Abcam, #ab420) antibody. The gels were run in 0.5x TBE with 10 mM MgCl_2_ for 3-5 hours at 90 V in an ice-water bath and imaged with a GE Image Want LAS 4000 imager (GE Healthcare). The gels with antibody-bound nanostructures were post-stained with 1x SYBR Gold (Thermo Fisher Scientific, #S11494) in the running buffer and destined overnight prior to imaging.

#### Cryo-Electron Microscopy

The structural characterization of the DNA nanostructures was done by cryo-electron microscopy (cryo-EM). A Vitrobot Mk4 (Thermo Fisher Scientific) was used to prepare cryo-EM specimens on Quantifoil R1.2/1.3 (LF, 300 Mesh, Copper) grids with an additional 2 nm carbon layer. The grids were cleaned using ozone cleaning for 2 min before applied to a 50 μL drop of 0.0001% (w/v) poly-L-lysine for 10 secs and allowed to air dry for 20 min. A total of 3 µl of the DNA origami solution was applied to the grid and allowed to equilibrate for 20 s before plunge freezing in liquid ethane. Data were collected with EPU software using a Krios G3i TEM operated at 300 kV. Images were acquired in 81 kx nanoprobe EFTEM SA mode with a slit width of 10 eV using a K3 Bioquantum. Exposure time was 4.3 s, during which 45 movie frames were collected with a fluency of 1.09 e Å−2 per frame. Image pre-processing until particle picking was carried out in Warp (Tegunov et al., 2019), with subsequent processing using CryoSPARC v4.5.3 (Punjani et al., 2017). An outline of the data processing pipeline can be found in **Figure S7**. For density visualization only, dust hiding on the reconstructed map was performed using UCSF ChimeraX (Pettersen et al., 2021).

#### Transmission Electron Microscopy

To visually inspect the DNA nanostructures, both bare and antigen-patterned with antibodies bound were negatively stained and observed by transmission electron microscopy (TEM). Carbon-coated Formvar grids (Electron Microscopy Sciences) were glow-discharged before applying 4 μl of 10-20 nM of DNA nanostructures for 20 seconds. The sample was blotted off on a filter paper, washed with MiliQ water for 4 s and stained for 20 s with 2% (w/v) aqueous solution of uranyl formate with 20 mM NaOH. The uranyl formate was also blotted off on a filter and the grids were air dried for about 45 min before imaging. Electron micrographs of the negatively stained samples were collected on a Talos 120C G2 operated at 120kV with a Ceta-D detector at 73 000x magnification in nanoprobe mode. Automated data collection was performed using the serialEM software suite to obtain individual micrographs consisting of a single frame recorded over a 2 sec exposure time with a fluence of 10eÅ^2^. Each of the datasets consisted of around 100 images, which were subsequently processed using CryoSPARC v4.5.3 (Punjani et al., 2017). Obtained reconstructions were visualized using UCSF ChimeraX (Pettersen et al., 2021). An outline of the data processing pipeline can be found in **Figure S6**.

#### Surface plasmon resonance

To compare with the PANMAP measurements, the binding kinetics of the digoxigenin-decorated nanostructures to the anti-digoxigenin antibodies were determined by surface plasmon resonance (SPR) using a Biacore T200 (GE Healthcare). Streptavidin was immobilized by amine coupling (Cytiva, #BR100050) on the surface of a Series S Sensor Chip CM3 (Cytiva, #BR100536) following the instructions from the manufacturer and using HBS-EP+ as running buffer (Cytiva, #BR100669) containing 10 mM HEPES, 150 mM NaCl, 3 mM EDTA, 0.05% v/v Surfactant P20 at pH 7.4. Briefly, the streptavidin was dissolved at a concentration of 40 μg/ml in 100 mM sodium acetate pH 4.5 and injected for 480 sec at a flow rate of 10 μl/min, obtaining around 900 RUs. Then, 100 nM of biotinylated oligonucleotides, complementary to the immobilization staples from the nanostructures, were injected for 120 sec at a flow rate of 5 μl/min followed by washing for 60 sec with the running buffer, yielding around 25 RUs of immobilized biotin-oligonucleotide. From this point on, all the injections and runs were done in running buffer HBS-EP+ supplemented with 5 mM MgCl_2_. The digoxigenin-decorated DNA nanostructures were diluted to 10 nM in the running buffer and injected for 400 s at a flow rate of 2 μl/min to immobilize about 500 RUs and washed for 60 s. In a parallel flow cell, empty nanostructures were immobilized to the same RU levels as a reference. To obtain the binding curves for analysis, single cycle kinetic runs were done with a 2-fold dilution of α-digoxigenin HRP antibody diluted from 192 nM to 12 nM. The contact time for each of the five concentrations was 500 s and the flow rate was 30 μl/min. After the last concentration, the dissociation phase was recorded for 1000 s and the surface was regenerated with two injections of 30 s with 50 mM NaOH. Representative sensorgrams of each step are shown in **Figure S9**. The sensorgram data were analyzed with the BIAevaluation 3.2 software (GE Healthcare) assuming a 1:1 Langmuir binding model. The whole run from the immobilization of the structures to the single cycle kinetics was repeated in three independent experiments.

#### qPCR

To experimentally determine the amount of DNA nanostructures immobilized on the wells for PANMAP, qPCR targeting the scaffold was performed. After the PANMAP experiment, the wells were washed once with milliQ water and airdried for a few minutes. To detach the DNA nanostructures from the plate, the wells were treated with 80 μl per well of 8 mM NaOH (Merck) and incubated for 15 minutes at room temperature. Then, the NaOH was diluted to 2.6 mM with water and 3 μl per well was collected for qPCR. The standard curve consisted of a 10-fold serial dilution of DNA origami structures, from 18.46 pM to 0.001846 pM, in the same NaOH buffer as the PANMAP samples. Using the Luna^®^ Universal qPCR Master Mix (NEB, #M3003) and following the instructions from the manufacturer, the qPCR reaction contained 10 μl of qPCR Master Mix, 0.5 μl of 10 μM for each primer targeting the scaffold (forward: CCAACGTGACCTATCCCATTAC, reverse: TTCCTGTAGCCAGCTTTCATC), 3 μl of target or standard curve sample, and up to 20 μl with nuclease-free water. The qPCR reaction was denatured for 1 minute at 95°C followed by amplification for 40 cycles of 15 seconds at 95°C and 30 seconds at 60°C using a StepOnePlus™ Real-Time PCR System (Thermo Fisher Scientific). As we aimed to test whether there was any difference in immobilization of the different origami structures, we performed a Kruskal Wallis test with a post-hoc Dunn’s test and the Benjamin Yekhouteli, adjusting for the false discovery rate which showed no statistically significant difference between the abundance of previously bound structures.

### PANMAP

The PANMAP experiments were performed in a strepatividin-coated 96-well plate (Thermo Fisher Scientific, #15126). First, the plate was washed three times with Wash Buffer containing 25 mM Tris, 150 mM NaCl, 0.1% BSA, 0.05% Tween and 5 mM MgCl_2_ using a volume of 200 μl per well, following the instructions from the manufacturer. For coating the surface, the library of decorated DNA nanostructures, including 1-antigen and 2-antigens separated by different distances, containing biotinylated anchor oligonucleotides, were diluted to 50 ng/μl of DNA in Wash Buffer. As a background and negative control, empty DNA nanostructures were always included. Each type of DNA nanostructure was immobilized in a different row of the plate with a volume of 50 μl per well and incubated for 2 hours at 23°C and 800 rpm. The unbound nanostructures were washed away three times with Wash Buffer using 200 μl per well. Then, the α-digoxigenin HRP (Abcam, #ab420) was prepared as a 5-fold dilution starting at 250 μg/ml and down to 0.016 μg/ml in Wash Buffer. A volume of 25 μl per well of antibody was allowed to bind to the antigen-DNA nanostructures for 2 hours at 37°C and 800 rpm, until reaching equilibrium. All the incubation steps were done using an Eppendorf ThermoMixer**™** C (Eppendorf, #5382000015) with a SmartBlock**™** Thermoblock for 96-well plates (Eppendorf, #5363000039) and a ThermoTop (Eppendorf, #5308000003) to prevent condensation. To avoid unbinding of the antibody during the washing process, the excess of antibody was removed well by well with an aspirator by adding and suctioning the Wash buffer three times. Finally, 100 μl per well of the HRP substrate 1-step**™** ABTS Solution (Thermo Fisher Scientific, #37615) was added to detect the bound antibodies. The HRP signal was measured as absorbance at 405 nm every 2 min over a period of 24 minutes using a Varioskan LUX plate reader (Thermo Fisher Scientific). The plate was shaken for 1 min and 50 s at 300 rpm and for 10 s at 900 rpm right before every absorbance measurement to avoid precipitation of the HRP product. The experiments were run in triplicates.

#### Conversion of PANMAP signal to structure occupancy

To convert the HRP absorbance signal into occupancy or mean number of antibodies bound per structure at steady state, we utilized the curve generated for the 1-antigen structure and the standard curve of moles of antibody bound vs absorbance unit.

For the 1-antigen nanostructure, the rate law expression for structures occupied by antibodies follows a 1-to-1 model:

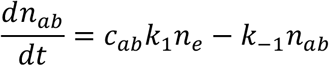

where *n*_*ab*_ is the number of structures occupied by antibodies, *n*_*e*_ is the number of unoccupied empty structures, *c*_*ab*_ is the solution phase concentration of antibody, and *k*_1_ and *k*_−1_ are the forward and reverse rate constants respectively.

At steady state, the equation is set to 0

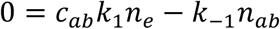

The number of unoccupied structures can be substituted

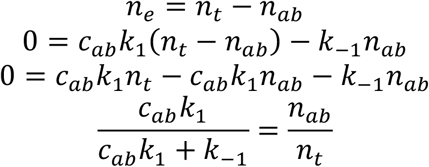

where *n*_*t*_ is the total number of structures.

The *K*_*D*1_ dissociation constant can be substituted to eliminate the reverse rate constant

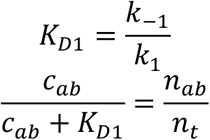

At saturation, for the 1-antigen sample, the total number of structures corresponds to the maximum number of structures occupied by an antibody 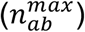

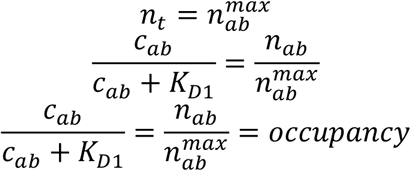

The equation for occupancy can be parameterized as linear function

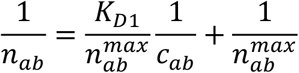

Where 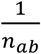 is the dependent variable, 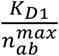 is the slope, 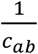 is the independent variable, and 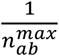 is the y-intercept. Like this, the *K*_*D*1_ and the 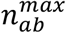 can be obtained from the linear form of equilibrium bound antibody where the inverse of the amount of bound antibody is plotted against the inverse of solution phase concentration of antibody.

#### Modeling and parameterizing steady state occupancy

To model the PANMAP signal, an expression was derived for the expected steady state structure occupancy or mean number of bound antibodies to a structure at steady state. The model is derived from a system of dynamic rate equations with the assumption that the rate of change of each state is equal to 0 at steady state.

For a 1 antigen system, the expected steady state occupancy follows a familiar 1-to-1 model:

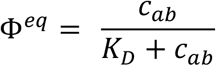

where *c*_*ab*_ is the solution phase concentration of antibody and *K*_*D*_ is the steady state dissociation constant of the antibody.

For the more complex, 2 antigen model, we follow a 5 state Markovian model in which an empty (State A) 2-antigen structure transitions to one of two monovalent states (States B and C) with a single antibody bound to one or other antigen sites, which may then transition to a bivalent state (State D) where a single antibody is bound to both antigens, or a saturated state (State E) where a second antibody binds to the other antigen. The model is based on a system of rate laws or ordinary differential equations for the rates of change 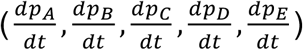 of the probabilities (*p*_*A*_, *p*_*B*_, *p*_*C*_, *p*_*D*_, *p*_*E*_) that a structure resides in a given state (A,B,C,D,E) of the 5 possible states, respectively.

The system of rate laws for the 5 states is:

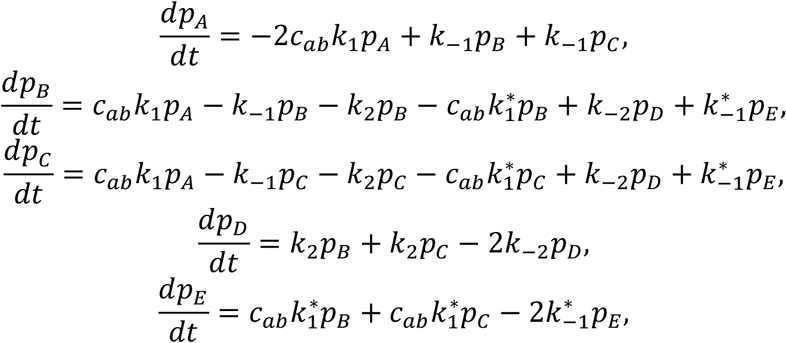

where *k*_1_ and *k*_−1_ are the monovalent forward and reverse binding rate constants, respectively, *k*_2_ and *k*_−2_ are the forward and reverse monovalent-to-bivalent transition rate constants respectively, and 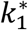 and 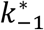 are the forward and reverse monovalent binding rate constants when binding is occurring under conditions of steric hindrance due to the presence of an already-bound monovalent antibody. This process is expected to be approach the behavior of the monovalent binding process when the separation distance between antigens is beyond the limit in which steric effects occur, whereas steric hindrance is expected to influence the process when the separation distance between antigens is below this critical distance.

The endpoint of the PANMAP experiment assumes steady state, or that the rates of change of the possible states are zero: 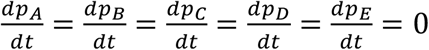.

Rearrangement of the steady state rate equation for State A, yields the familiar dissociation constant:

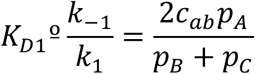

The steady state rate equation for State D, may be rearranged to yield a dissociation constant for the monovalent-to-bivalent transition process, a function of separation distance x between antigens:

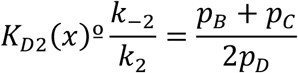

Finally, the steady state rate equation for State E may be rearranged to yield a dissociation constant for the distance-dependent, sterically hindered, monovalent binding when an adjacent monovalent antibody is already present prior to binding:

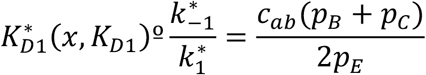

An expression for the mean structure occupancy can be derived by first deriving the steady state probabilities for each state in terms of the the dissociation constants. The dissociation constants substitution enables 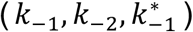 to be eliminated, yielding steady state equations in terms of the forward rate constants and the dissociation constants:

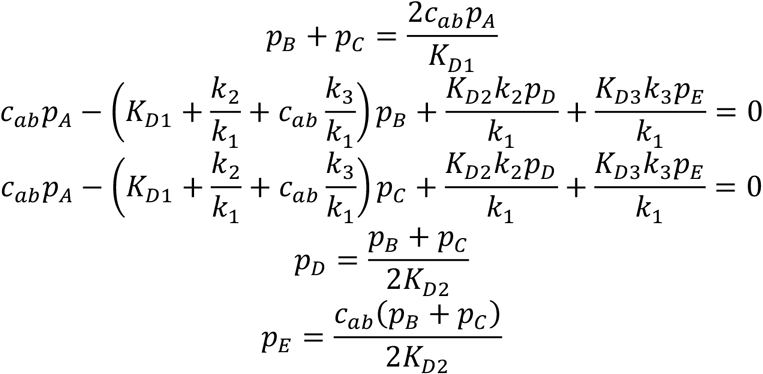

The normalization condition applies to the state probabilities, i.e. that the sum of all state probabilities must sum to 1:

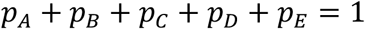

By substituting from previous equations, it is possible to express the normalization condition formula to be only in terms of *p*_*A*_, *K*_*D*1_, *K*_*D*2_, 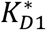, *c*_*ab*_

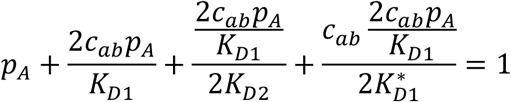

Which can be arranged to solve for *p*_*A*_

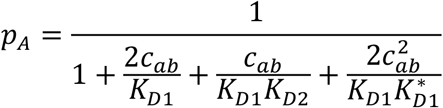

The remaining probabilities can be expressed in terms of *p*_*A*_ :

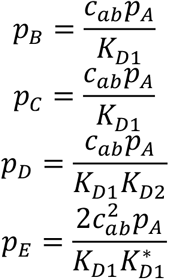

Substitution with these equations then allows each of the probabilities to be expressed in terms of solution concentration and the dissociation constants only. After simplification the following equations are obtained:

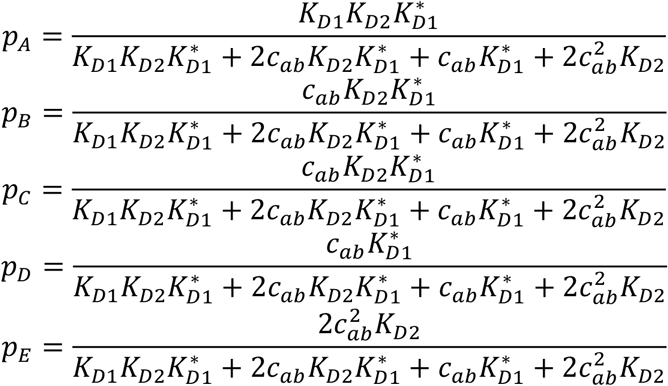

Expressions for steady state probability of each system state may then be combined with knowledge of the occupancy of each state, i.e. the number of antibodies bound to a structure in each state.

For the empty state A, the occupancy is

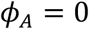

For the single-antibody monovalent and bivalent states B,C, and D, the occupancy is

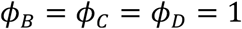

And for the saturated state E, the occupancy is

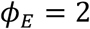

The expected occupancy value is then obtained via a summation of occupancy values of each possible state weighted by the probabilities of the respective states:

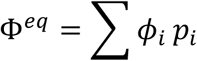

By substituting the formulae for the state probabilities into the expected occupancy equation, we arrive at an expression for the steady state occupancy that is in terms of the dissociation constants and the solution phase concentration.

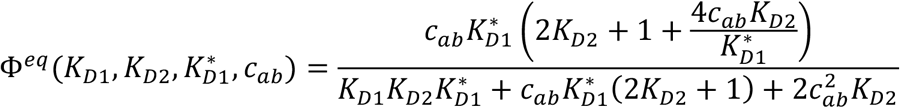

To parameterize the model of steady state occupancy, the occupancy signal data from a PANMAP experiment in which solution phase antibody concentration was varied is used as the basis for fitting the above expression for expected occupancy, changing the dissociation constant variables to minimize the loss ℒ:

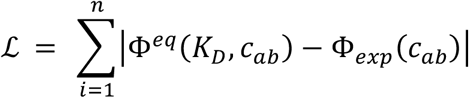

for the 1-antigen structure, and

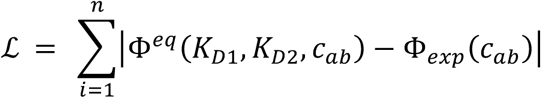

in the 2-antigen structures, where Φ_*exp*_(*c*_*ab*_) is the experimental occupancy signal from the PANMAP data.

#### Modeling distance dependence of dissociation constants

To model the dissociation constants as a function of distance, phenomenological equations were conceived to model the distance-dependent monovalent-to-bivalent transition, represented by the parameter *K*_*D*2_, and the sterically hindered monovalent binding, represented by the parameter 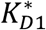.

A symmetric parabolic function of separation distance x was chosen to model the trade-off behavior of the monovalent-to-bivalent transition, in which weak binding was observed at both close and far separation distances, while strong binding was observed for intermediate distances.

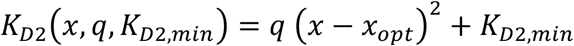

where *q* is a parameter representing the spatial tolerance of the antibody, or its ability to dynamically stretch and compress to accommodate different antigen separation distances, and *K*_*D*2,*min*_ is the minimum dissociation constant at the bivalent optimum or distance in which bivalent binding is maximally favored.

To model the dependence of the monovalent binding to an unoccupied antigen adjacent to a monovalently occupied antigen, an exponential decay model was used to capture the decrease in dissociation constant observed as a function of separation distance, with larger distances approaching the unhindered low (strong) monovalent dissociation constant observed for unoccupied monovalent binding.

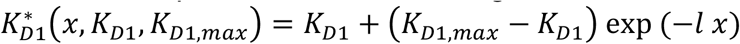

where l is a physical parameter representing influence of steric hindrance on the monovalent binding dissociation, and *K*_*D*1,*max*_ is the dissociation constant at x=0 separation distance. Here, *K*_*D*1_ is a static parameter which is unaffected by antigen separation distance when it comes to the initial binding onto an unoccupied empty structure.

#### Modeling mixed distance ELISA signal

To model the signal of a randomly coated surface of antigens, e.g. for classical ELISA, in which antigens have variable distances between one another, it is possible to combine the expression for the expected occupancy as a function of distance with the commonly known function for the nearest neighbor distance probability distribution. The expected occupancy, as a function of the Poisson density *λ* (antigens/unit area), is:

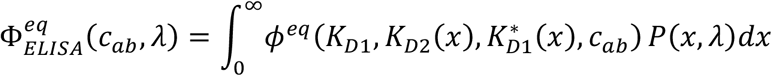

where *P*(*x, λ*) is the probability distribution of nearest neighbor inter-antigen distances:

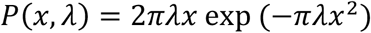

Here it must be noted that this model does not account for >2 body interactions, which will result in a greater contribution to steric hindrance than predicted by the model.

### Statistics

Data are reported as mean ± S.E.M.

## Code availability

All the code used to produce the results of this study including installation, demonstration and result reproduction instructions are available on GitHub (https://github.com/molecular-programming-group/PANMAP-processing).

## Acknowledgments

Electron microscopy data was collected at the Karolinska Institutet 3D-EM facility (https://ki.se/cmb/3d-em).

## Authors contribution

IRL: Conceptualization, methodology, formal analysis, validation, investigation, writing-original draft preparation, editing, visualization.

IB: Validation, formal analysis, investigation, editing, visualization.

SKD: Methodology, validation, formal analysis, editing, visualization.

ITH: Conceptualization, methodology, formal analysis, validation, writing -original draft preparation, editing, supervision, visualization.

BH: Supervision, writing, reviewing, editing, project administration, funding acquisition.

