## Supplementary figures for "Resolving antibody avidity through nanoscale antigen patterning"

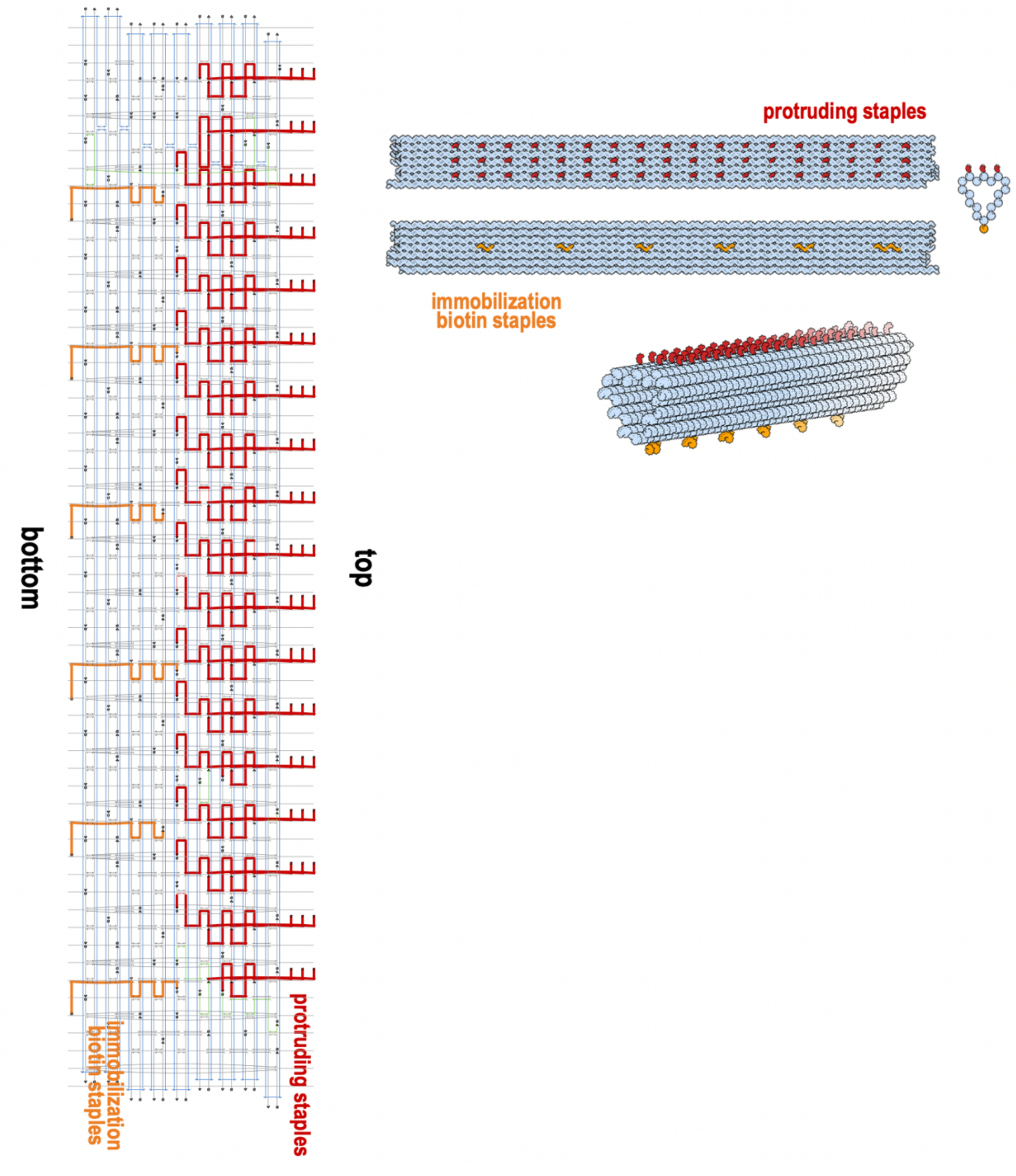


**Figure S1. DNA origami rod scaffold routing and staple design.** The DNA origami rod was composed of 18 DNA helices arranged in a honeycomb lattice. Protruding staples on the top surface were used for the anchoring of antigens, and protruding staples on the bottom surface for biotin-streptavidin immobilization of the structures on the assay plate.

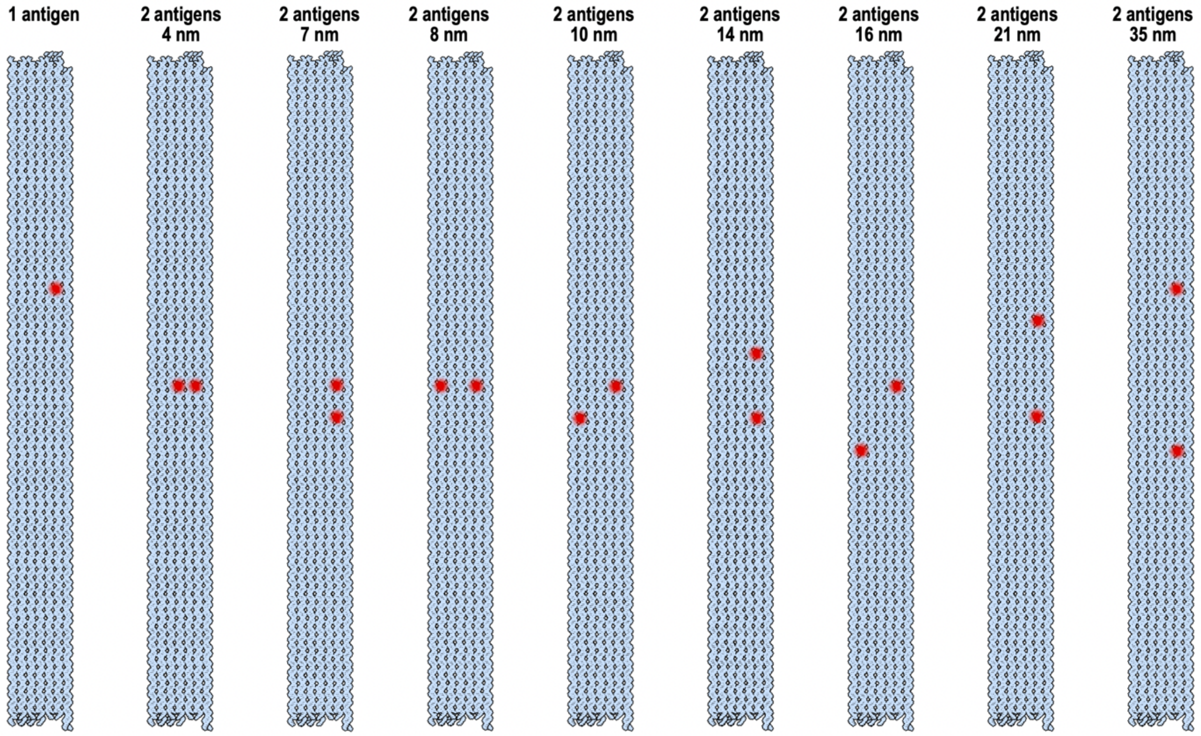


**Figure S2. Library of protruding staples combinations for 1 antigen or 2 antigens separated at different distances on the rod origami surface.**

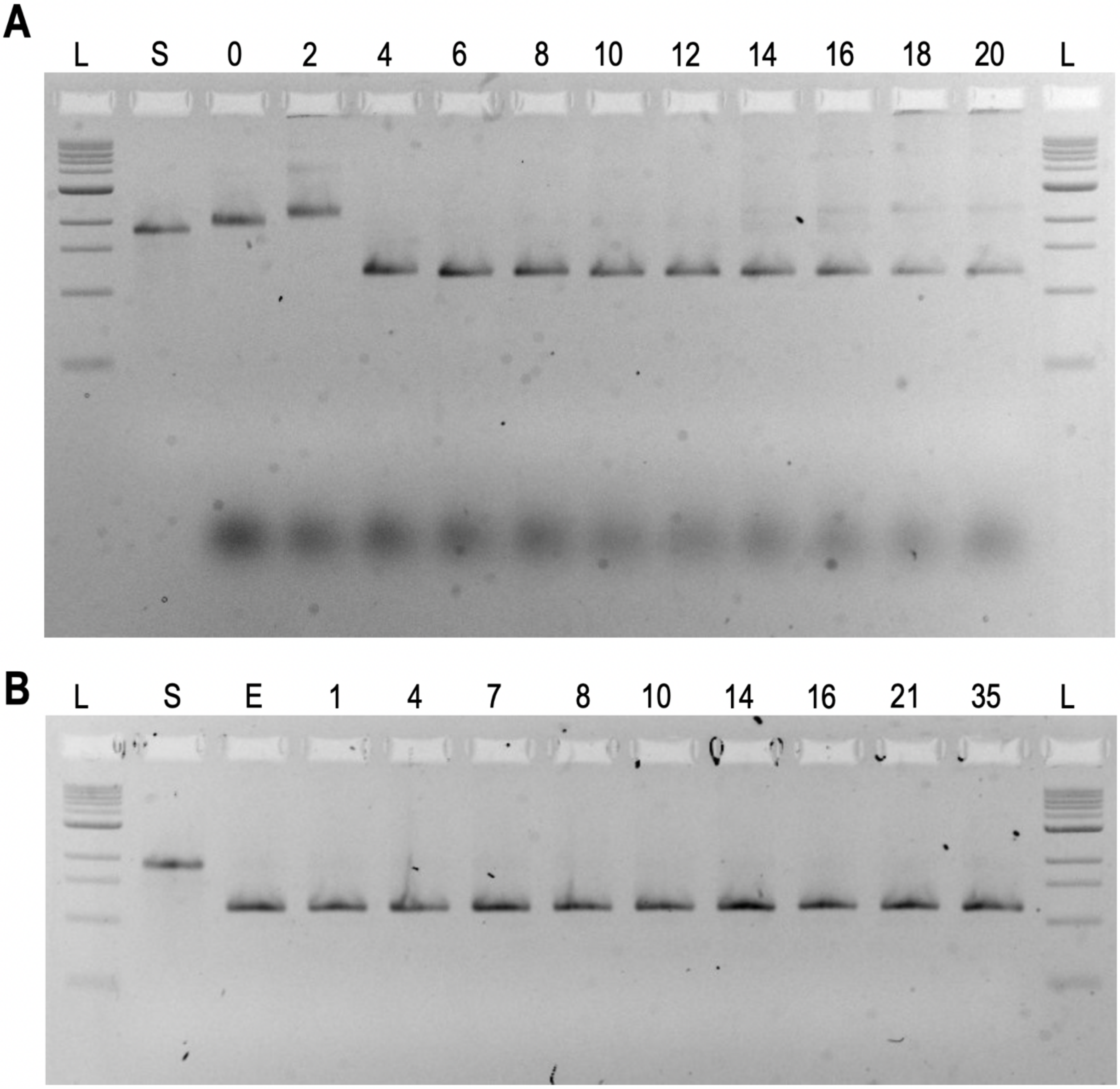


**Figure S3. Agarose gels of the rod DNA origami. (a)** EMSA of magnesium screening for the rod origami. L: DNA ladder, S: p7560 scaffold, 0-20: increasing amount of $MgCl_2$ from 0 mM to 20 mM. 2% agarose gel with ethidium bromide. **(b)** EMSA of digoxigenin-patterned rod origami after folding and PEG precipitation. L: DNA ladder, S: p7560 scaffold, E: empty origami, 1: 1-antigen digoxigenin origami, 4-35: 2-antigen origami separated by increasing distances from 4 nm to 35 nm. 2% agarose gel with ethidium bromide.

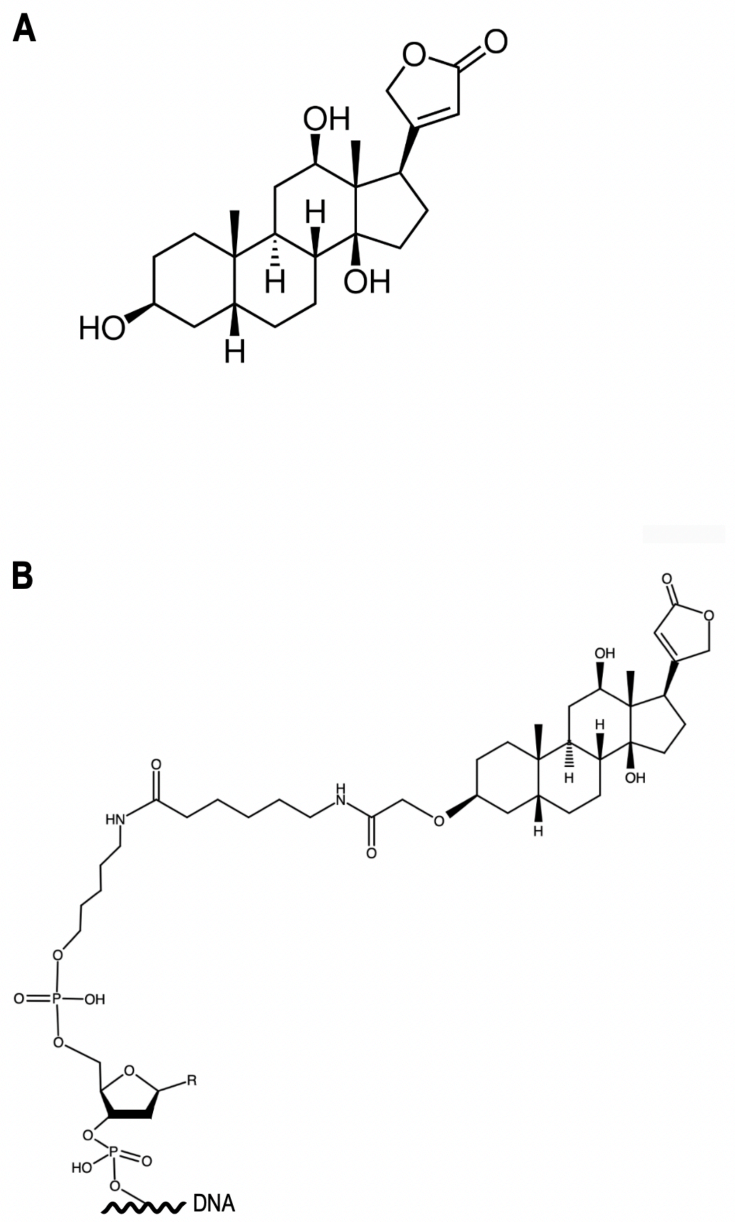


**Figure S4. Chemical structure of the digoxigenin hapten (A) and of 5’ digoxigenin modification for the protruding staples (B).**

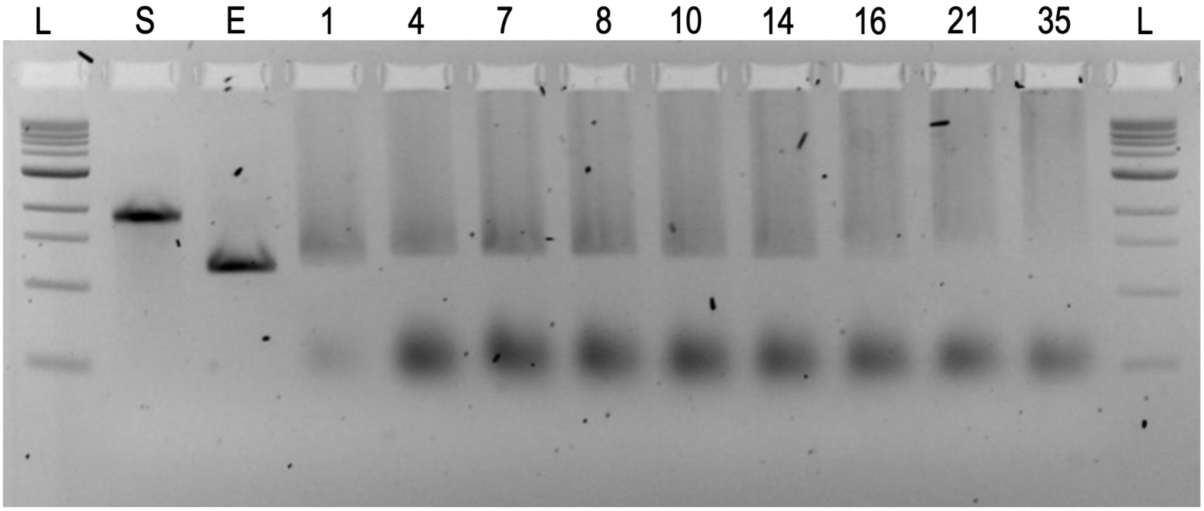


**Figure S5. Agarose gel of digoxigenin-pattern rod origami with bound anti-digoxigenin antibody.** EMSA of digoxigenin-origami after incubation with a 100-fold molar excess of HRP anti-digoxigenin antibody for 1 hour at 37°C. L: DNA ladder, S: p7560 scaffold, E: empty origami, 1: 1-digoxigenin origami with antibody, 4-35: 2-digoxigenin origami separated by distances from 4 to 35 nm with antibody. 2% agarose gel with ethidium bromide.

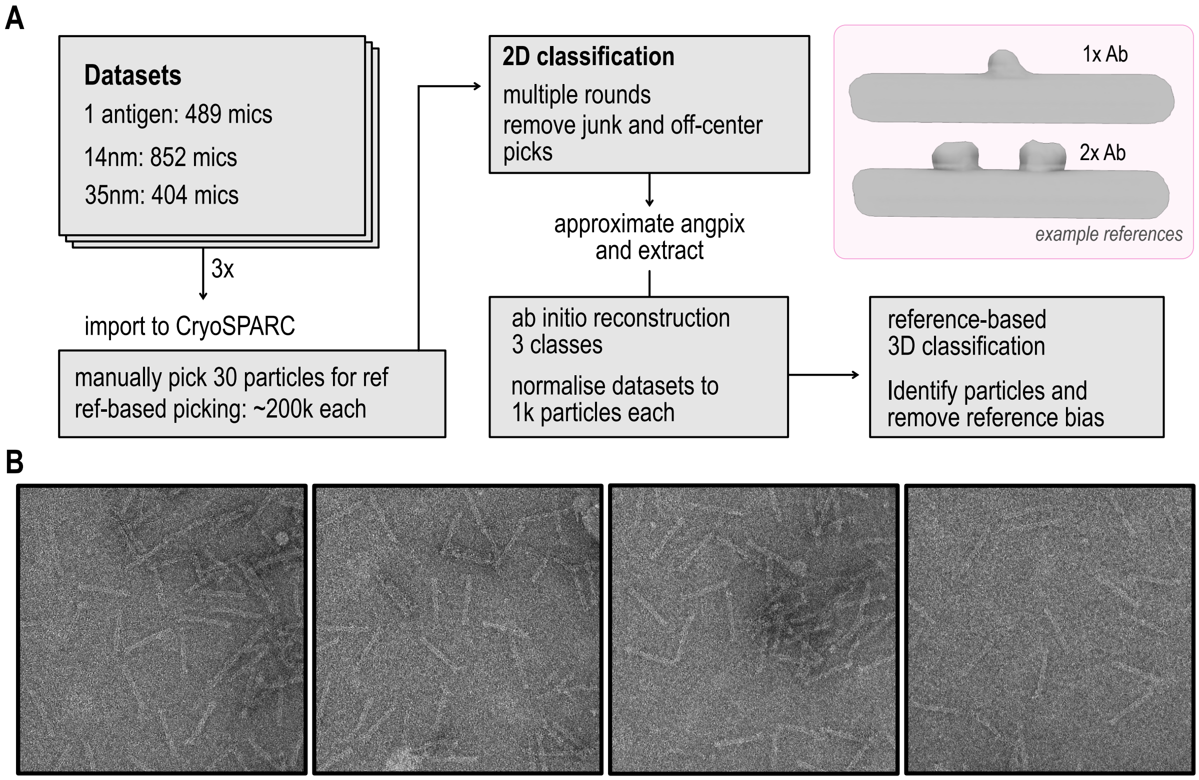


**Figure S6. Outline describing the processing pipeline used during reconstruction of the three rod DNA origami nanostructures containing either 1-antigen, 2-antigens at 14 nm or 2-antigens at 35 nm after NS-TEM imaging. (A)** Data processing was normalised among the three datasets. Example references used were generated via applying the molmap function in ChimeraX with a value of 100. Reference using one antibody (1x Ab) was applied for processing datasets with a single antigen and a 14nm spacer, while the reference using two antibodies (2x Ab) was applied for processing the 35nm spacer dataset. **(B)** Representative micrographs from the datasets.

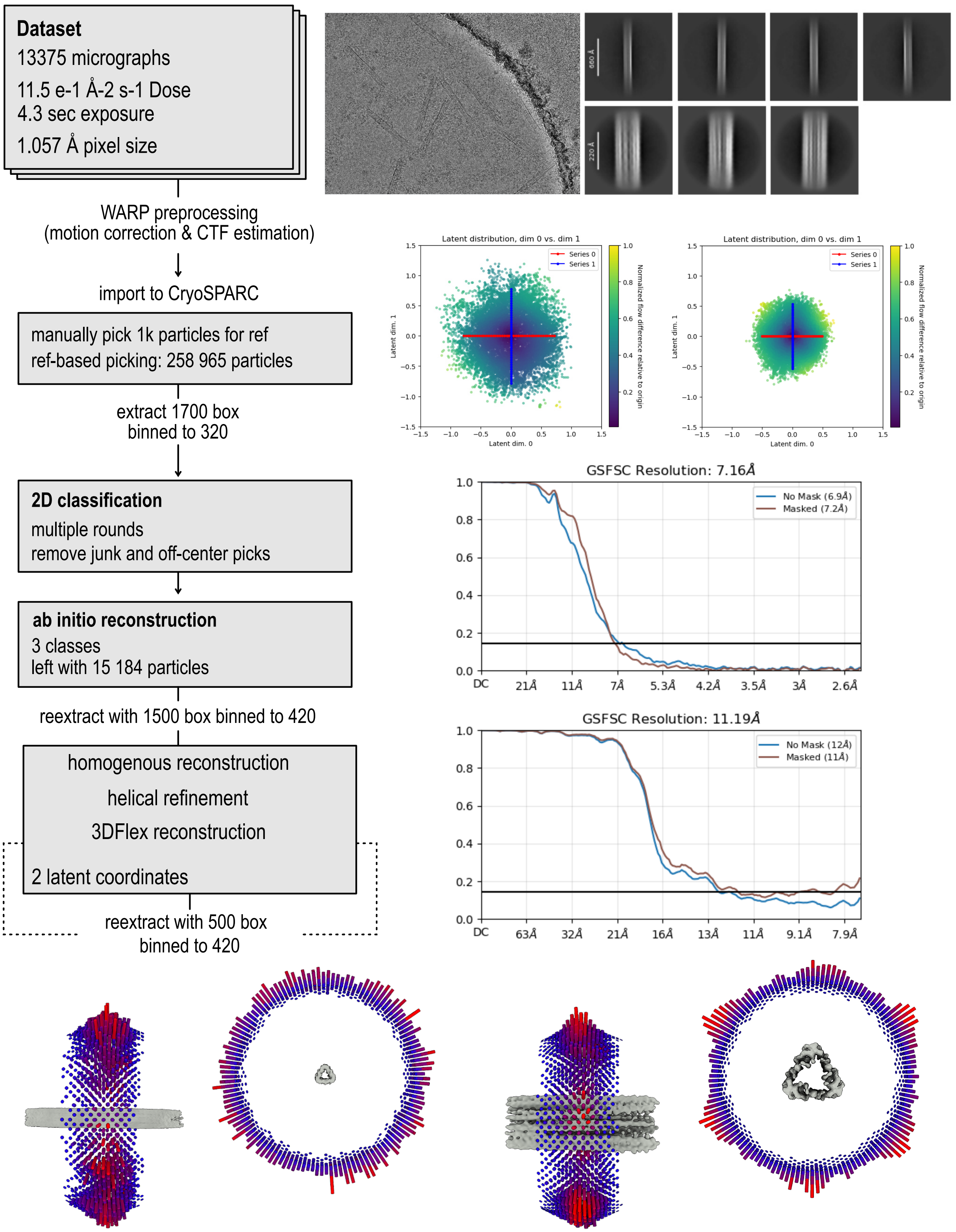


**Figure S7. Outline describing the preprocessing and processing steps to obtaining the final Cryo-EM reconstruction of the DNA origami rod.** The outline includes the processing pipeline; a representative micrograph of the datasets; 2D classes of both full-lenght and core reconstructions; the latent distribution plots obtained during 3DFlex analysis for both full-lenght and core reconstructions; FSC curves from the 3DFlex reconstruction for both the full-lenght and core reconstructions; and the angular view distribution for both the full-lenght and core reconstructions.

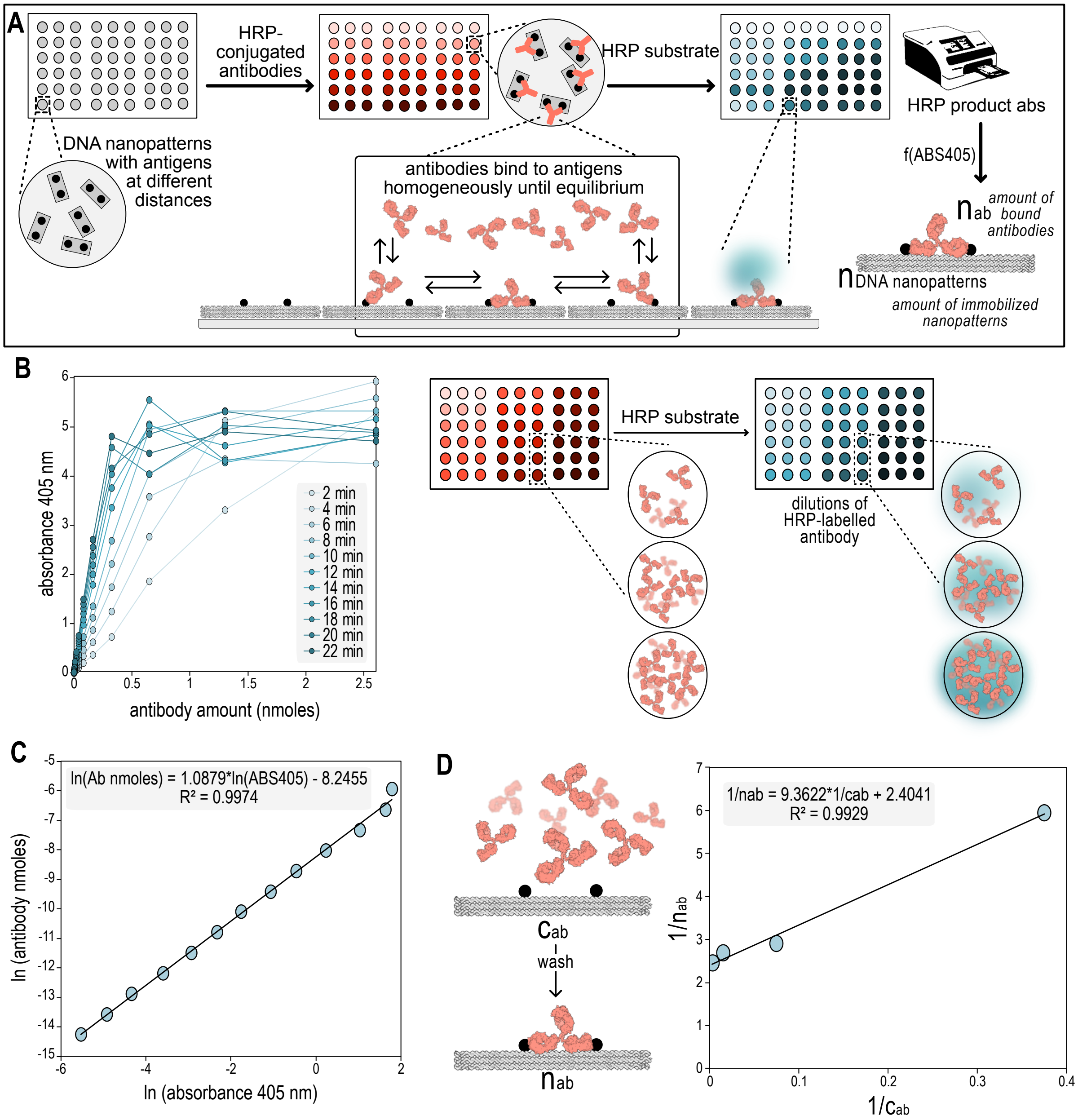


**Figure S8. Pipeline to convert HRP absorbance to absolute quantity of DNA nanopatterns and bound antibodies. (A)** Experimental workflow for a PANMAP experiment. **(B)** Experimental standard curves of the HRP product absorbance (y-axis) from different HRP-labelled antibody amounts (x-axis) measured after different incubation timepoints (grey inset). **(C)** HRP conversion factor corresponding to the linearized standard curve of a 10-minutes incubation obtained in (B). The function of the curve and the coefficient of determination (R) are shown in the grey inset. **(D)** Curve representing the relationship between bound antibodies per structure (nab), and amount of antibodies in solution (Cab) for the 1-antigen DNA nanopattern after converting the HRP units with the conversion factor from (C).

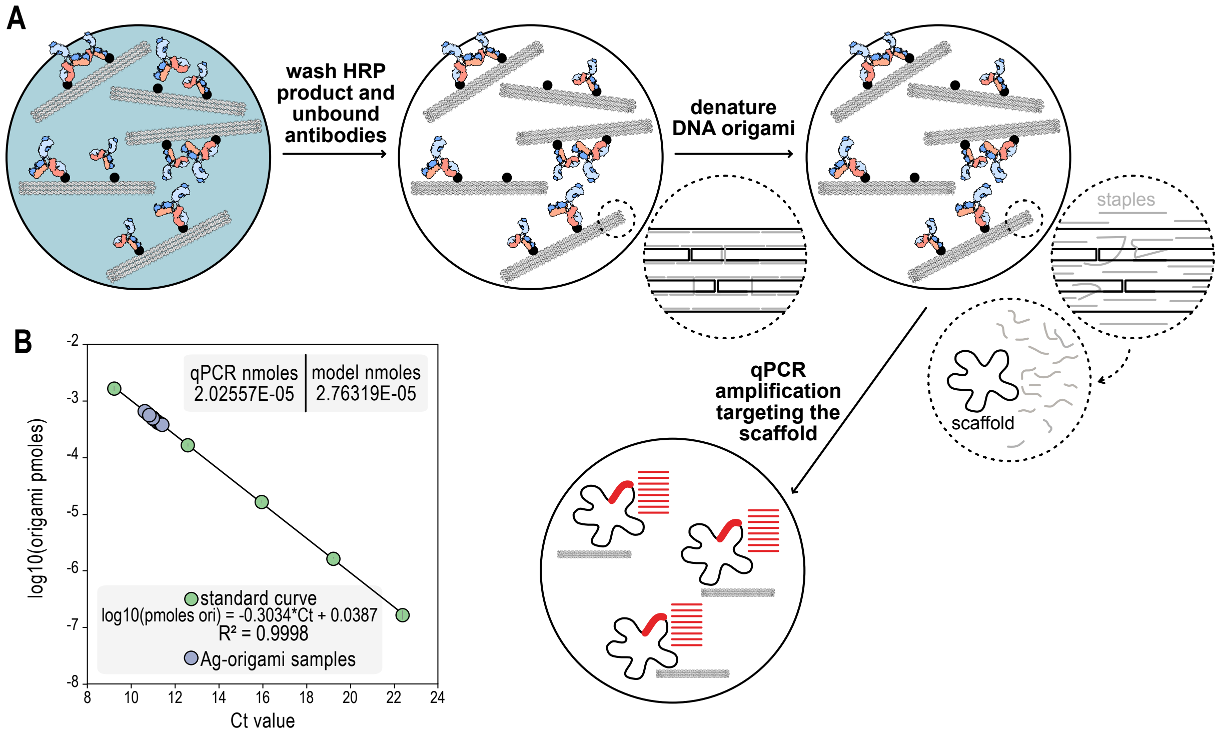


**Figure S9. qPCR to validate the absolute quantity of DNA nanostructures obtained from the HRP conversion factor. (A)** Schematic of the protocol to measure the absolute quantity of DNA nanostructures by qPCR amplification targeting the scaffold. **(B)** Standard curve obtained after qPCR amplification of the scaffold (green dots), and the random DNA nanopattern samples quantified with the same primers (purple dots) to verify the homogeneity of structure immobilization in PANMAP. The function of the standard curve and the coefficient of determination (R) are shown in the top grey inset. The calculated amount of nanostructures using the curve in Figure S8D and the quantified amount of structures using qPCR are shown in the bottom grey box.

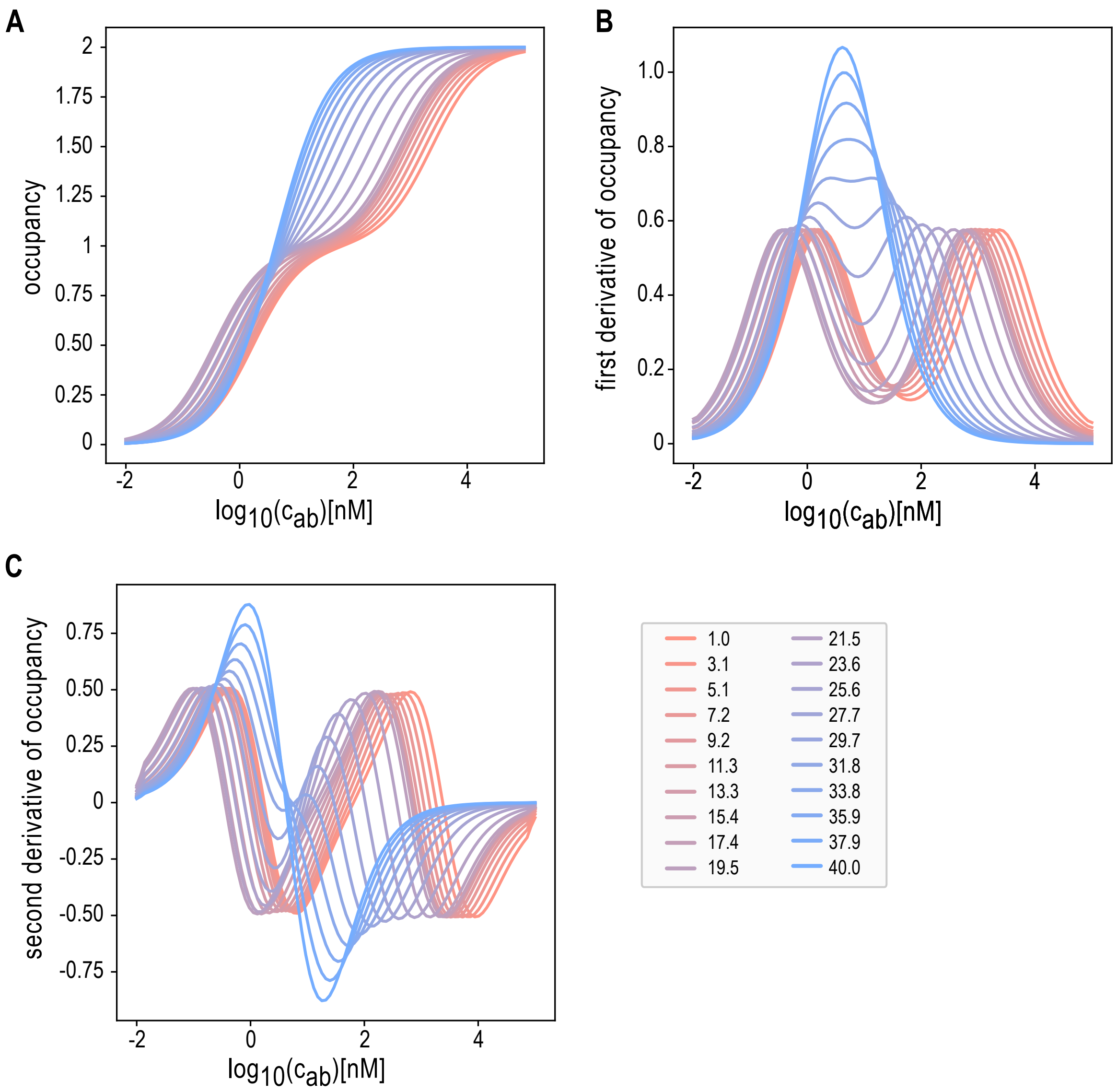


**Figure S10. Distance-dependent binding behaviour revealed by occupancy curves and their derivatives. (A)** Steady-state occupancy calculated with the 3 KD model plotted against log antibody concentration for antigen separations 1 - 40 nm. At short separations the curves are double-sigmoidal, exhibiting a convex shoulder that signals a dominant bivalent state. As the separation grows, the shoulder diminishes and by 30-40 nm the curves become single-sigmoidal, consistent with monovalent binding. **(B)** First derivative of steady state occupancy highlighting the regime switch. Near optimal spacings <15 nm show two distinct maxima: the low concentration peak marks the monovalent to bivalent transition, while the high concentration peak marks the bivalent to saturated binding condition. With increasing distance, the peaks translate leftward (lower concentration) from short to intermediate separations, indicating enhanced avidity, then coalesce into a single peak at large separations where bivalent binding is lost. **(C)** Second derivative of steady state occupancy maps the curvature of the binding curves. Zero-crossings correspond to the inflection points seen in (A), while their merger into a single crossings at large x mirrors the peak coalescence in (B). The alternation of positive and negative components encodes both the emergence and eventual disappearance of the bivalent shoulder as antigen spacing increases.

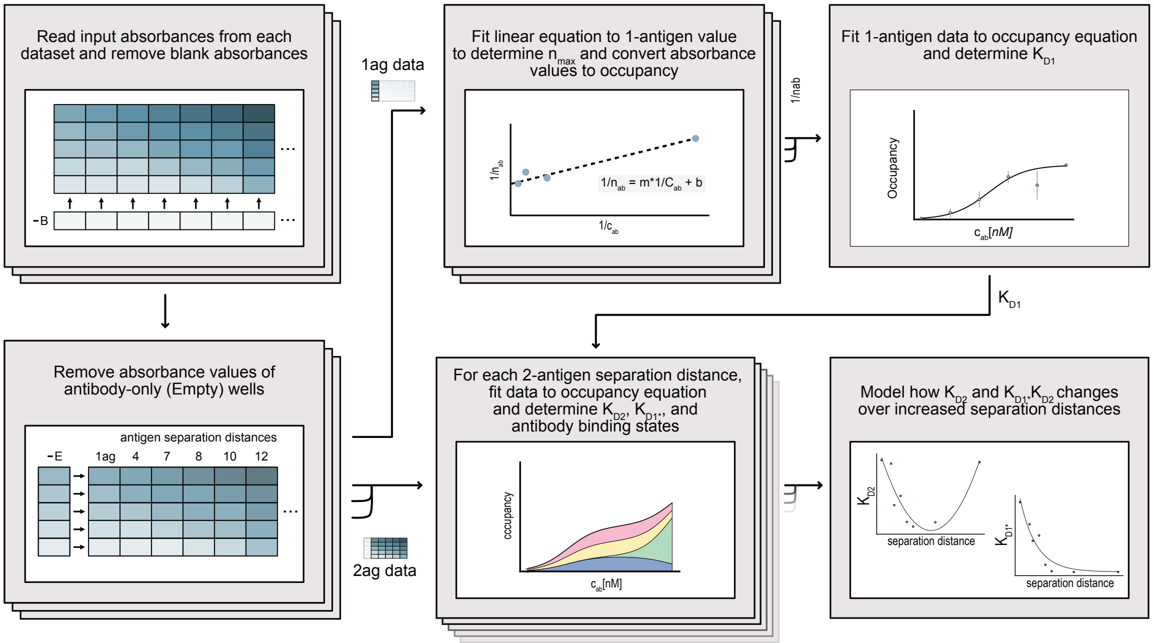


**Figure S11. Pipeline for processing replicate PANMAP data.** First, each sample is corrected by substraction of buffer-only wells (blank) and construct-free wells (empty). Then, $n_{max}$ is calculated by using the 1-antigen data together with an antibody amount to HRP signal standard curve. $n_{max}$ is then used together with the 1-antigen data to determine $K_{D1}$, which, together with the 2-antigen data, is the input to model $K_{D2}$, $K_{D1*}$, and antibody binding state ratios as a function of antigen separation distances.
